# A reabsorption mechanism in *Drosophila* renal tubules maintains amino acid homeostasis

**DOI:** 10.64898/2026.05.07.723391

**Authors:** Ayano Oi, Pattama Wiriyasermkul, Hiroto Hakamada, Ayako Isomura-Matoba, Miyu Kubota, Dooseon Cho, Osamu Nishimura, Yusuke Kato, Chisako Sakuma, Takefumi Kondo, Shushi Nagamori, Masayuki Miura, Fumiaki Obata

**Affiliations:** Laboratory for Nutritional Biology, RIKEN Center for Biosystems Dynamics Research, Kobe, Hyogo, 650-0047, Japan; Department of Food and Agricultural Sciences, Iwate University, Morioka, Iwate, 020-8550, Japan; Department of Biological Chemistry and Food Science, Iwate University, Morioka, Iwate, 020-8550, Japan; Center for Systemic Intelligence in Biomedicine, The Jikei University School of Medicine, Tokyo, 105-8461, Japan; Laboratory of Molecular Cell Biology and Development, Graduate School of Biostudies, Kyoto University, Sakyo-ku, Kyoto, 606-8501, Japan; Genomics Research and Analysis Support Team, RIKEN Center for Biosystems Dynamics Research, Kobe, Hyogo, 650-0047, Japan; Laboratory for Developmental Genome System, RIKEN Center for Biosystems Dynamics Research, Kobe, Hyogo, 650-0047, Japan; Metabolic and Behavioral Physiology RIKEN ECL Research Team, RIKEN Center for Biosystems Dynamics Research, Kobe, Hyogo, 650-0047, Japan; Metabolic and Behavioral Physiology RIKEN ECL Research Team, RIKEN Pioneering Research Institute, Kobe, Hyogo, 650-0047, Japan; Division of Multiscale Nutritional Biochemistry, Research Center for Medical Sciences, The Jikei University School of Medicine, Tokyo, 105-8461, Japan; Laboratory for Cell Vigor Regulation, National Institute for Basic Biology, Okazaki, Aichi, 444-8585, Japan

## Abstract

The kidney plays a central role in maintaining systemic homeostasis by regulating the excretion of metabolic compounds. In mammals, this is achieved through filtration followed by selective reabsorption, whereas insect Malpighian tubules (MTs) are thought to function primarily through active secretion. Although reabsorption of ions and water has been described in MTs, whether these tissues regulate metabolite balance through reabsorption remains unclear. Here, we uncover an amino acid (AA) reabsorption mechanism in *Drosophila* MTs. Quantification of AAs in excreta reveals that AA excretion changes rapidly and markedly in response to dietary protein levels, while systemic AA levels remain stable. Using *ex vivo* assays, we show that MTs actively transport AAs. We identify the AA transporter Slimfast (Slif), expressed in the lower tubule, as a key mediator required to maintain systemic levels of basic AAs. These findings reveal a previously unrecognised reabsorption-based strategy in insect renal tubules that functionally parallels mammalian kidney physiology, despite their distinct organisation. Our study demonstrates that AA excretion forms an additional layer to achieve metabolic homeostasis.

## Introduction

Excretion plays a crucial role in maintaining homeostasis by controlling the quality of blood. Mammalian kidney, the organ responsible for urine production, is composed of functional units called nephrons, which consist of the glomerulus and renal tubules. Urine is generated from primary filtration of low-molecular-weight substances at the glomerulus, followed by the reabsorption and secretion in the renal tubules. This two-step process is essential for the efficient elimination of waste products while retaining essential ones from circulation. Among these, amino acids (AAs) are indispensable nutrients for living organisms and are therefore believed to be almost completely reabsorbed in the proximal tubules^1^.

The insect excretory system is composed of nephrocytes, Malpighian tubules (MTs), and the hindgut^2–4^. Nephrocytes are specialised filtration cells that express genes homologous to those in mammalian podocytes in glomerulus^5^. However, unlike in mammals, nephrocytes are not physically connected to MTs, and thus haemolymph filtration by nephrocytes does not contribute to urine formation. MTs, the insect renal tubules, are connected to the junction between the midgut and hindgut and maintain osmotic homeostasis by producing urine^2,3^. In the main segment of MTs, ions and water secretion from the haemolymph is driven by a proton gradient generated by V-ATPase, and mediated by various channels and transporters in both principal and stellate cells^6^. The primary urine is further processed by reabsorption from tubular lumen into the haemolymph in the lower tubules, the proximal region of MTs, as well as in the hindgut^3,7^. Ion and water transport mechanisms in these excretory organs have been extensively studied and are thought to rely primarily on active secretion^6,8^. In contrast, although the possibility of AA transport in insect MTs was suggested decades ago^9^, the descriptive and mechanistic investigation of this process remain poorly conducted. In particular, whether and how AAs are reabsorbed has not been directly demonstrated.

AAs require membrane transporters for their distribution, as they cannot freely diffuse across lipid bilayers. In mammals, more than 60 solute carrier (SLC) transporters mediate AA transport^10^, many of which play essential roles in renal reabsorption. Although homologous transporters are present in insects^11^, their physiological roles—particularly in excretory organs—are still largely unknown. Among insect AA transporters, Slimfast (Slif), the *Drosophila* ortholog of SLC7A1 (CAT1), has been implicated in AA sensing and metabolic homeostasis. *slif* was originally identified as a regulator of body size in *Drosophila* larvae, where it promotes organismal growth by importing AAs into fat body cells, thereby activating mTORC1 signaling and stimulating the secretion of insulin-like peptides^12–14^. In addition, Slif-mediated AA sensing in both the fat body and dopaminergic neurons contributes to the regulation of feeding behaviour^15,16^. In addition, Slif can contribute to nutrient absorption in the gut, where its knockdown extends and its overexpression shortens lifespan^17^. However, its function in MTs, as well as its substrate specificity and physiological roles in excretory processes, remain largely unknown.

Here, we investigate the mechanisms and physiological significance of AA handling in *Drosophila* excretory organs. By integrating dietary manipulation, genetic approaches, and quantitative metabolite analysis, we uncover an unexpected layer of regulation in AA excretion. We demonstrate that the conserved SLC transporter Slif mediates selective AA reabsorption in the MTs, thereby contributing to systemic AA homeostasis, especially for basic AAs such as arginine (Arg) and histidine (His). These findings challenge the conventional view that insect MTs rely predominantly on selective secretion and instead reveal that reabsorption-based control of metabolite excretion is an evolutionarily conserved strategy.

## Results

### Amino acid excretion changes in response to dietary protein levels

To scrutinise the impact of protein intake, adult wild-type Canton-S (CS) flies were fed with various concentrations (1, 2, 4, 6, 8, and 12%) of yeast extract (YE), as a major protein source. Lifespan decreased as YE concentration increased in both females and males (Fig. 1a, b). In females, lifespan was dose-dependent with the longest at 1% YE and the shortest at 12% YE. Fecundity after one-day feeding of YE diet increased in a YE concentration dependent manner (Fig. 1c), indicating a trade-off between lifespan and reproduction^18,19^. However, after seven days of feeding, the fecundity declined at 12% YE, suggesting a toxicity associated with prolonged high protein intake (Fig. 1d). In another wild-type strain, Dahomey, 12% YE diet shortened lifespan but increased fecundity (Extended Data Fig. 1a–c). Based on these results, we defined 12% YE as a high-protein condition for subsequent experiments.

**Fig. 1.**
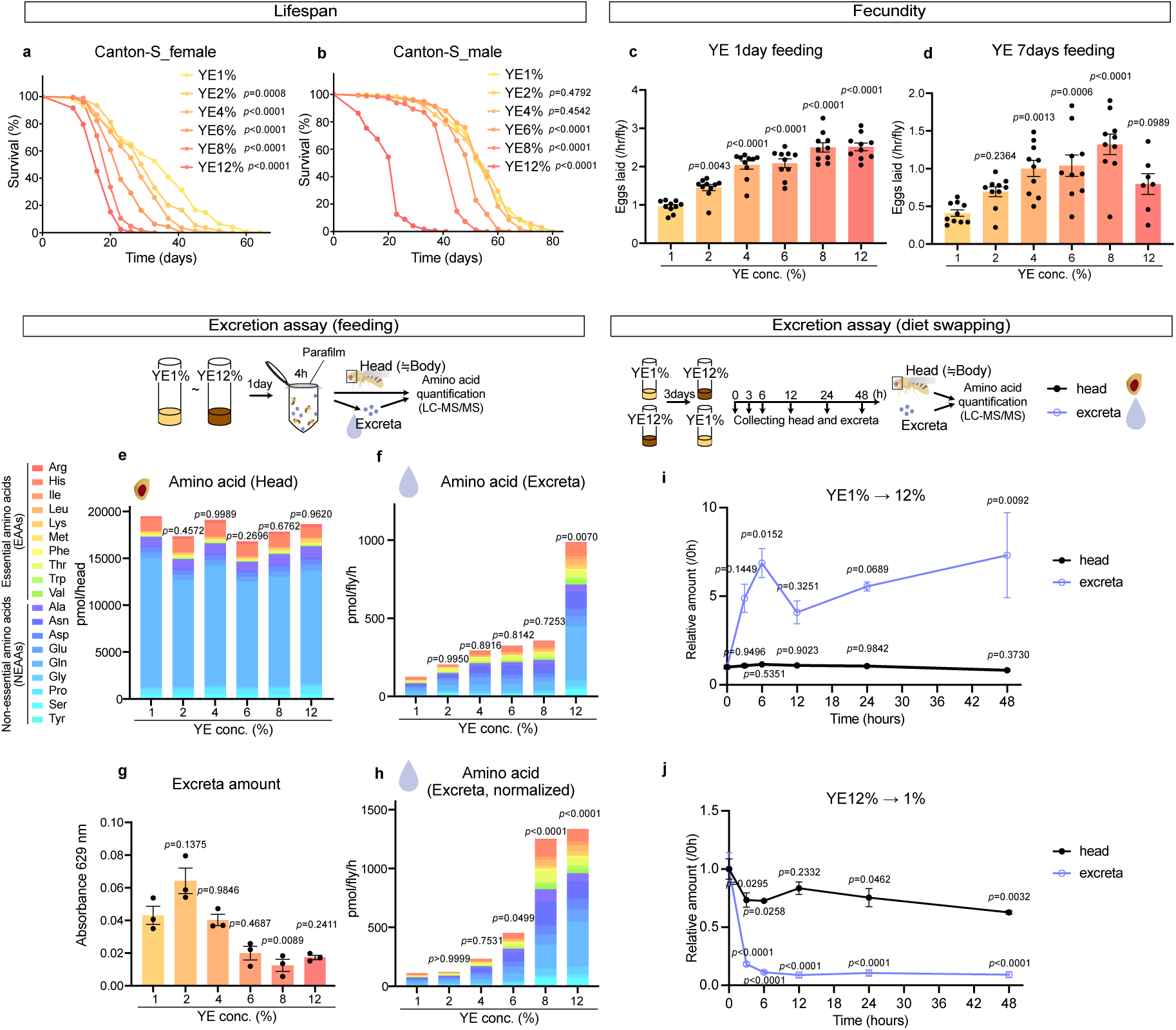
Amino acid excretion depends on dietary yeast levels. **a, b** Lifespan of female (**a**) and male (**b**) Canton-S (CS) flies reared on a diet containing yeast extract (YE) at indicated concentrations. n=169 for YE 1% female, n=165 for YE 2% female, n=171 for YE 4% female, n=158 for YE 6% female, n=170 for YE 8% female, n=166 for YE 12% female, n=171 for YE 1% male, n=176 for YE 2% male, n=174 for YE 4% male, n=175 for YE 6% male, n=178 for YE 8% male, and n=175 for YE 12% male. Statistics: log-rank test. **c, d** Fecundity of female CS flies reared on diet containing YE at indicated concentrations for 1 day (**c**) or 7 days (**d**). n=7 for 12% 7 days condition and n=10 for all other conditions. Statistics: one-way ANOVA with Šidák’s multiple comparisons. **e, f, h** Quantification of AAs in heads (**e**), excreta (**f**), and excreta normalised to total excreta amount (**h**) in female CS flies after feeding on diets containing the indicated concentrations of YE for 1 day, followed by 4 h incubation in tubes. n=3. Statistical analysis was performed using the total AA levels. Statistics: one-way ANOVA with Šidák’s multiple comparisons. **g** Absorbance at 629 nm of excreta remaining in the tubes and solubilised in buffer. n=3. Statistics: one-way ANOVA with Šidák’s multiple comparisons. **i, j** Relative amounts of total AAs in heads (black) and excreta (blue) of female CS flies, normalised to 0 h, after switching the diet from 1% YE to 12% YE (**i**) or from 12% YE to 1% YE (**j**). At the indicated time points, flies were placed in tubes for 2 h to collect excreta. Statistical analysis was performed by comparing each time point with 0 h. n=3. Statistics: one-way ANOVA with Šidák’s multiple comparisons. Each graph shows the mean ± SEM.

We then examined the excretion of AAs in response to dietary protein intake. We quantified AA levels in body parts and excreta based on a previously described excretion assay^20^. Female flies were fed YE diets supplemented with a blue dye for one day to visualise excreta. The group of four flies were allowed to excrete in a 1.5 mL tubes for 4 hours. Heads and excreta were collected and subjected to AA analysis by LC-MS/MS. The head contains the brain, haemolymph, and fat body, enabling systemic metabolite measurements without potential impacts of gut contents and ovaries (including eggs). Total AA levels in the head remained relatively stable despite varied YE concentration (Fig. 1e). However, individual AAs showed distinct response. While many essential AAs (EAAs) increased with higher YE concentrations, His, threonine (Thr), and many non-essential AAs (NEAAs) remained relatively constant (Extended Data Fig. 2). This is partly consistent with previous observation reporting that His, as well as Arg, levels are maintained even under low or high protein diet^21,22^. Internal tyrosine (Tyr) levels increased significantly at 8% and 12% YE compared with 1% YE (Extended Data Fig. 2), exceptionally among NEAAs. This is relevant to the fact that Tyr is a nutritionally maintained AA which predominantly regulates nutrient sensing machineries in *Drosophila*^21,23^. In contrast to the head samples, the total amount of AAs in the excreta correlated well with the dietary intake and showed a marked elevation between 8% and 12% YE (Fig. 1f). Consistent with the total AA profile, many individual AAs were markedly increased at 12% YE (Extended Data Fig. 3). As an indicator of total excretion volume, we quantified the blue dye amount in the excreta. Dye levels tended to decrease as YE concentration increased from 2% to 8% (Fig. 1g). Because excretion volume has been reported to reflect food intake^24^, reduced feeding within this range may contribute to preventing excess in internal AA levels. When normalised AA amounts to blue dye levels, we found that flies fed high protein diets, particularly 8% and 12% YE, excreted higher amounts of AAs (Fig. 1h), suggesting that AAs in excreta are concentrated under these dietary conditions. We initially speculated that the AA composition in excreta might reflect their abundance within the body. However, normalising excreted AA amounts to their respective body levels revealed a reduced contribution of NEAAs, whereas EAAs such as isoleucine (Ile), leucine (Leu), tryptophan (Trp), and valine (Val) each exhibited excreta-to-head ratios exceeding 0.2 in 12% YE condition (Extended Data Fig. 4a). Interestingly, AAs are relatively evenly represented in YE^21^, and this composition differs from those observed in both the head and the excreta (Extended Data Fig. 4b). These results suggest that dietary AAs are not simply excreted in proportion to intake but are instead subject to selective absorption and excretion. Different AAs may be regulated by distinct homeostatic mechanisms.

Because YE contains various nutrients in addition to AAs and proteins, we next used a fully synthetic diet, or holidic medium^25,26^, to specifically manipulate dietary AA content. AA levels were adjusted to one-quarter (0.25×) or 3-fold (3×) relative to the standard diet (1×). Similar to the YE manipulation, increased dietary AAs shortened lifespan of both females and males (Extended Data Fig. 5a, b). Fecundity was decreased upon one day of feeding with the 3× AA diet (Extended Data Fig. 5c), likely reflecting high AA stress, consistent with the decline observed after prolonged feeding with 12% YE (Fig. 1d). Although body AA levels also increased with higher dietary AA levels, AA excretion was markedly elevated in both females and males (Extended Data Fig. 5d, e). An inverse correlation between dietary AA concentration and total excretion volume was observed particularly in females (Extended Data Fig. 5f). Overall, changes in dietary AA alone recapitulated the impact of YE manipulation on AA excretion, with even more pronounced effects. Because AAs in the synthetic diet are present in free form, their bioavailability should be higher than that of protein-bound AAs in YE, which could explain the stronger phenotypes observed.

To determine how rapidly changes in AA excretion occur after dietary shifts, we switched YE concentrations from 1% to 12% or from 12% to 1% and measured AA levels in bodies and excreta over 48 hours. When YE was increased from 1% to 12%, internal AA levels remained largely unchanged, whereas AA levels in the excreta increased about 7-fold within 6 hours after swapping the diet (Fig. 1i). Conversely, when YE was reduced from 12% to 1%, internal AA levels again showed minimal change, but excreted AAs decreased to approximately one-fifth of the initial level within 3 hours (Fig. 1j). These results suggest that pronounced changes in excretion following dietary YE shifts contribute to maintaining internal AA homeostasis.

### Amino acids injected into haemolymph are excreted

It was possible that dietary AAs pass through the gut and were excreted without absorption. To negate this possibility, we directly injected AAs into the haemolymph. Using the AA composition of the synthetic diet^25^, a 4-fold concentrated AA cocktail dissolved in PBS was injected into the haemolymph through a glass capillary. Body AA levels were measured before or after injection, and the excreta were collected for 4h (Fig. 2a). Immediately after injection, internal AA levels were increased to approximately 1.6-fold in the AA-injected group, which was decreased over the following 4 hours (Fig. 2b). These results indicate that injected AAs were metabolised and/or excreted. Notably, EAA levels returned to baseline in 4 hours of AA injection (Fig. 2c), suggesting that excretion plays a prominent role in regulating internal EAA levels. Glutamine (Gln) levels slightly increased after injection (Fig. 2b), consistent with the possibility that excess AAs are metabolised into Gln. Expectedly, total AAs in excreta were markedly increased following AA injection compared to the control PBS injection (Fig. 2d), indicating that AAs injected into the haemolymph was eliminated via urinary excretion. To estimate the contribution of excretion to the decrease in body EAAs over 4 hours, we normalised the units and calculated the reduction in each EAA during this period. The proportion of excreted EAAs relative to the total decrease was highest for His (34.4%), followed by Trp (13.3%), whereas that of all EAAs was 6.2% (Fig. 2e). These results indicate that excretion contributes to a greater extent to the regulation of His levels compared to other AAs, whereas other AAs are more likely to be metabolised. Although *Drosophila* possesses histidine decarboxylase and can convert His into histamine, it lacks histidine ammonia-lyase and therefore cannot metabolise His into glutamate (Glu) or Gln (KEGG Pathway^27^). This limited metabolic capacity may account for the high proportion of His being excreted unchanged.

**Fig. 2.**
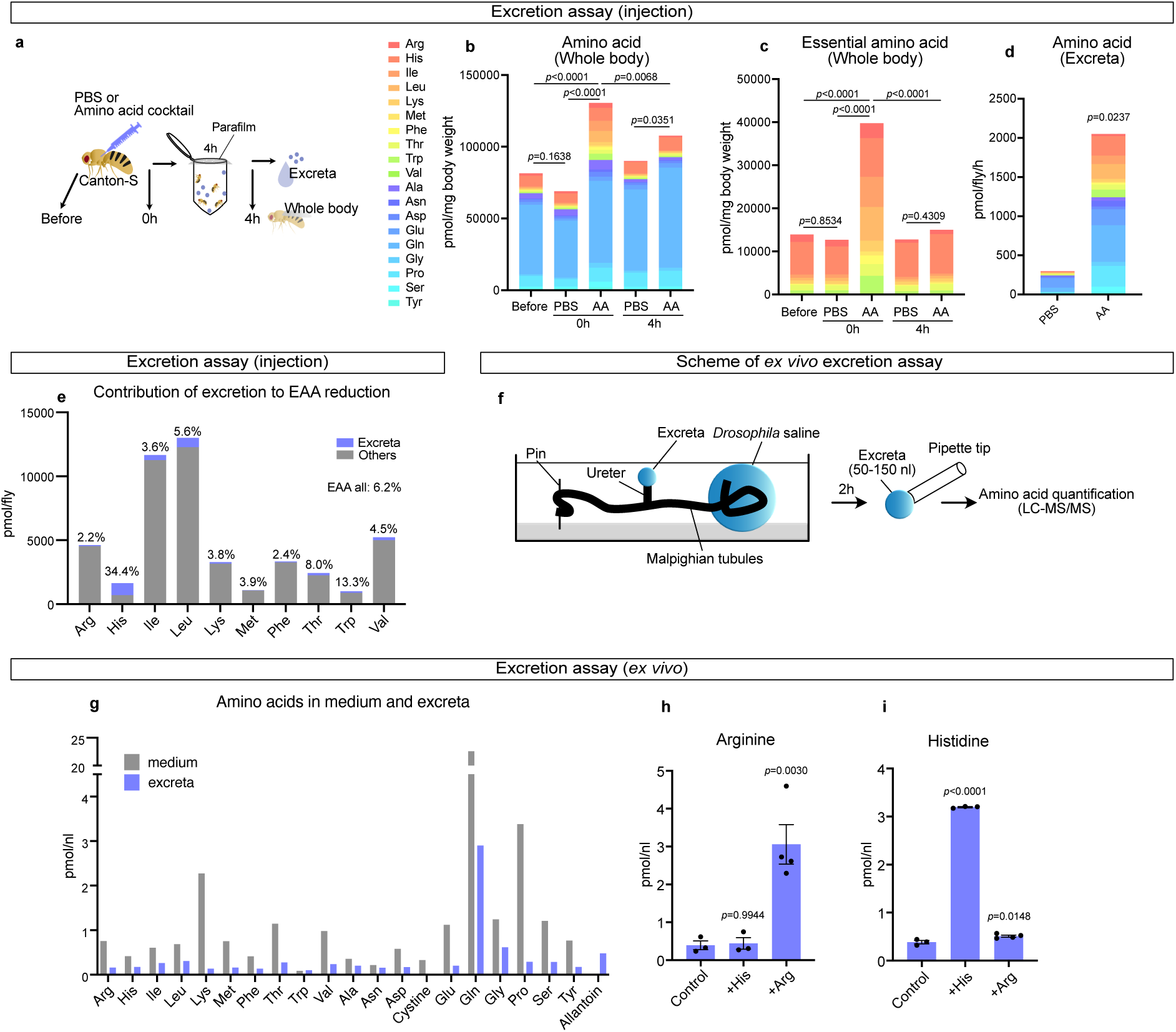
Amino acids are excreted from the haemolymph via the Malpighian tubules. **a** Schematic of the excretion assay following injection. **b, c** Quantification of all AAs (**b**) and EAAs (**c**) in whole bodies of female CS flies before and after injection with PBS (control) or an AA cocktail. n=3. Statistical analysis was performed using the total AA levels. Statistics: one-way ANOVA with Tukey’s multiple comparisons. **d** Quantification of AAs in the excreta of female CS flies 4 h after injection with PBS (control) or an AA cocktail. n=3. Statistical analysis was performed using the total levels. Statistics: unpaired two-tailed Student’s t test. **e** The proportion of excreted EAAs relative to the total decrease in the body for each AA. Values above the bars indicate the fraction of the decrease accounted for by excretion for each EAA. **f** Schematic of the *ex vivo* excretion assay. **g** Quantification of AAs in the medium (gray) and the excreta (blue) of female CS flies collected during the *ex vivo* excretion assay. n=1. **h, i** Quantification of Arg (**h**) and His (**i**) in excreta of female CS flies collected during the *ex vivo* excretion assay with medium supplemented with 10 mM Arg (**h**) or 10 mM His (**i**). n=4 for 10mM Arg, and n=3 for all other conditions. Statistics: one-way ANOVA with Šidák’s multiple comparisons. Each graph shows the mean ± SEM.

To further visualise the excretion of exogenous AAs, we used a stable isotope labelled His ([¹³C₆,¹⁵N₃]L-His) and distinguished injected His from endogenous pools. Labelled His levels in the body increased immediately after injection and showed a decreasing trend in both males and females after 3 hours (Extended Data Fig. 6a, b). Because the injected solution contained a much higher concentration of [¹³C₆,¹⁵N₃]L-His (100 mM) than the His present in the 4× AA cocktail (16.9 mM), the decline in body levels likely occurred more slowly than cocktail injection. Interestingly, [¹³C₆,¹⁵N₃]L-His accounted for approximately 48.4% of total excreta in females and 58.0% in males (Extended Data Fig. 6c), indicating that excretion is not simply the elimination of newly acquired AAs.

### Malpighian tubules regulate amino acid excretion

Although AAs in haemolymph can be excreted via MTs, the mechanisms of AA transport remain largely unknown. To directly determine whether the MTs have the capacity to transport AAs, we performed *ex vivo* excretion assays, based on the Ramsay assay^28–30^. Secreted fluid from isolated tubules in a dish was collected with a fine pipette tip and analysed by LC-MS/MS (Fig. 2f). Notably, allantoin, the final product of purine metabolism, which is generated from uric acid in MTs^20^, was also detected in the secreted fluid (Fig. 2g), confirming that the collected droplets represented tubule derived secretion. We detected almost all AAs in excreta, indicating that MTs possess a route for AA transport (Fig. 2g). Because cysteine (Cys) could not be reliably measured due to its susceptibility to oxidation, cystine was measured instead. Many AAs were excreted in proportions similar to their abundance in the medium (average 35.1%), including Gln, which was highly abundant in both. In contrast, lysine (Lys) was excreted at a markedly lower proportion (5.9%), possibly reflecting its metabolism in MTs, where the degradation enzyme LKRSDH is expressed (FlyAtlas2 Anatomy RNA-seq^31^). Furthermore, supplementation of the culture medium with a single AA, such as Arg or His, increased the excretion of the corresponding AAs (Fig. 2h, i). Together, these results demonstrate that MTs can directly transport AAs.

### Homeostatic control of excretion in response to dietary AA levels

Our data suggested that flies have ability to regulate AA excretion in response to changes in internal AA levels. However, it remained unclear whether this regulatory mechanism also works at the level of individual AAs, and if so, how flies determine the threshold to or not to excrete AAs. To address this, we manipulated single AA levels using a synthetic diet by focusing on His, whose internal levels appear to be more strongly regulated by excretion compared to other AAs (Fig. 2e). We prepared diets with various His concentrations from 0× (depletion) to 1× (4.2 mM) of the standard synthetic diet. After one week of feeding, internal His levels increased in response to dietary His intake (Extended Data Fig. 7a). In contrast, excreted His levels remained low when dietary His was below 0.4× (Extended Data Fig. 7b). When dietary His exceeded 0.6×, the flies begun to excrete His (Extended Data Fig. 7b), which was further upregulated upon 1×. To examine the physiological response to changes in dietary His, we measured lifespan and fecundity. Lifespan was slightly shortened at 0× and 10× His but showed little change at intermediate concentrations (Extended Data Fig. 7c), consistent with the previous report that His deficiency has relatively modest effects on lifespan among EAAs^32^. In contrast, fecundity was very low at 0× His and reached plateau when dietary His was 0.4× or higher (Extended Data Fig. 7d). Taken together, flies do not excrete His until its internal levels are beyond the threshold of maximising fecundity, which might be a proxy of anabolic reaction, indicating the mechanism of regulating excretion based on physiological demand. Therefore, monitoring excreted His may provide a practical way to estimate the dietary requirement for His. It is noteworthy that internal His levels continued to increase even at dietary concentrations above 0.6×, suggesting that the elevation of His cannot be fully compensated by excretion.

### Manipulation of a single amino acid affects excretion of other amino acids

Next, we tried to investigate whether manipulation of a single AA influences the excretion of other AAs. To test this, we supplemented a 4% YE diet with either His or Lys to level equivalent to those in a 12% YE diet. One day after feeding the His supplemented diet, excretion of basic AAs, including Arg, His, and Lys, increased, together with several EAAs such as Ile, Leu, Thr, Trp, and Val (Extended Data Fig. 7e). In contrast, Lys supplementation increased the excretion of Arg, His, and Lys (Extended Data Fig. 7f). On the other hand, depletion of His from the diet reduced the excretion of Arg and Lys in addition to His (Extended Data Fig. 7g), although these changes were not fully reflected in whole body AA levels (Extended Data Fig. 7h). To examine the excrete capacity of AA from the haemolymph, flies were pre-fed a 0× His or 0× Lys diet for one week and then injected with an AA cocktail containing both His and Lys. Under His depletion, excretion of Arg and Lys tended to decrease together with His (Extended Data Fig. 7i). In contrast, when flies were pre-fed a 0× Lys diet, excretion of Lys and Arg, but not His, showed a decreasing trend (Extended Data Fig. 7j). Taken together, these results suggest that there are several transport pathways shared among subsets of AAs (several EAAs, basic AAs, and Lys/Arg), and that their capacity might be regulated depending on the dietary AA levels.

To describe potential transcriptional changes in AA transporters in response to His or Lys deficiency, we performed transcriptome analysis of the MTs from flies fed 0× His or 0× Lys diets for three days. Approximately 70% of the genes significantly altered under His 0×, and 50% of those altered under Lys 0× were shared between the two conditions (Extended Data Fig. 8a, b, Table S1). Among the commonly upregulated genes, those involved in ribosome biogenesis was strongly enriched, whereas commonly downregulated genes were enriched for functions related to catabolic processes (Extended Data Fig. 8a, b, Table S1). Many putative AA transporters were similarly regulated under both conditions. However, the expression level of several transporters, including *CG1607*, *List*, and *CG16700* exhibited specific changes depending on the depleted AA (Extended Data Fig. 8c), suggesting that these transporters may regulate distinct excretion pattern.

### An amino acid transporter Slimfast localised in the lower tubule in MTs

Next, we dissected the AA transport mechanism in MTs. We focused on Slif since it is the most well-characterised AA transporter in *Drosophila*, whose function in the MTs has not been examined. MTs consist of multiple cell types and are divided into functionally distinct regions (Fig. 3a). The main segment, which occupies the largest area, generates primary urine by secreting ions and water from the haemolymph into the lumen^2^. The lower tubule and the ureter reabsorb ions and water from the lumen to the haemolymph to regulate osmorality^7^. These secretion and reabsorption are mainly reported in the context of ions and water, and the transport of metabolites including AAs in MTs are not fully understood. To determine the expression pattern of *slif* in MTs, we first investigated the single-cell transcriptome datasets (DRSC RNA Seq explorer^33^,https://www.flyrnai.org/tools/rna_seq/web/). *slif* showed relatively higher expression in principal cells in the lower tubule and the ureter (Extended Data Fig. 9a), a region known to reabsorb ions and water^7^, suggesting that Slif may reabsorb AAs into the haemolymph.

**Fig. 3.**
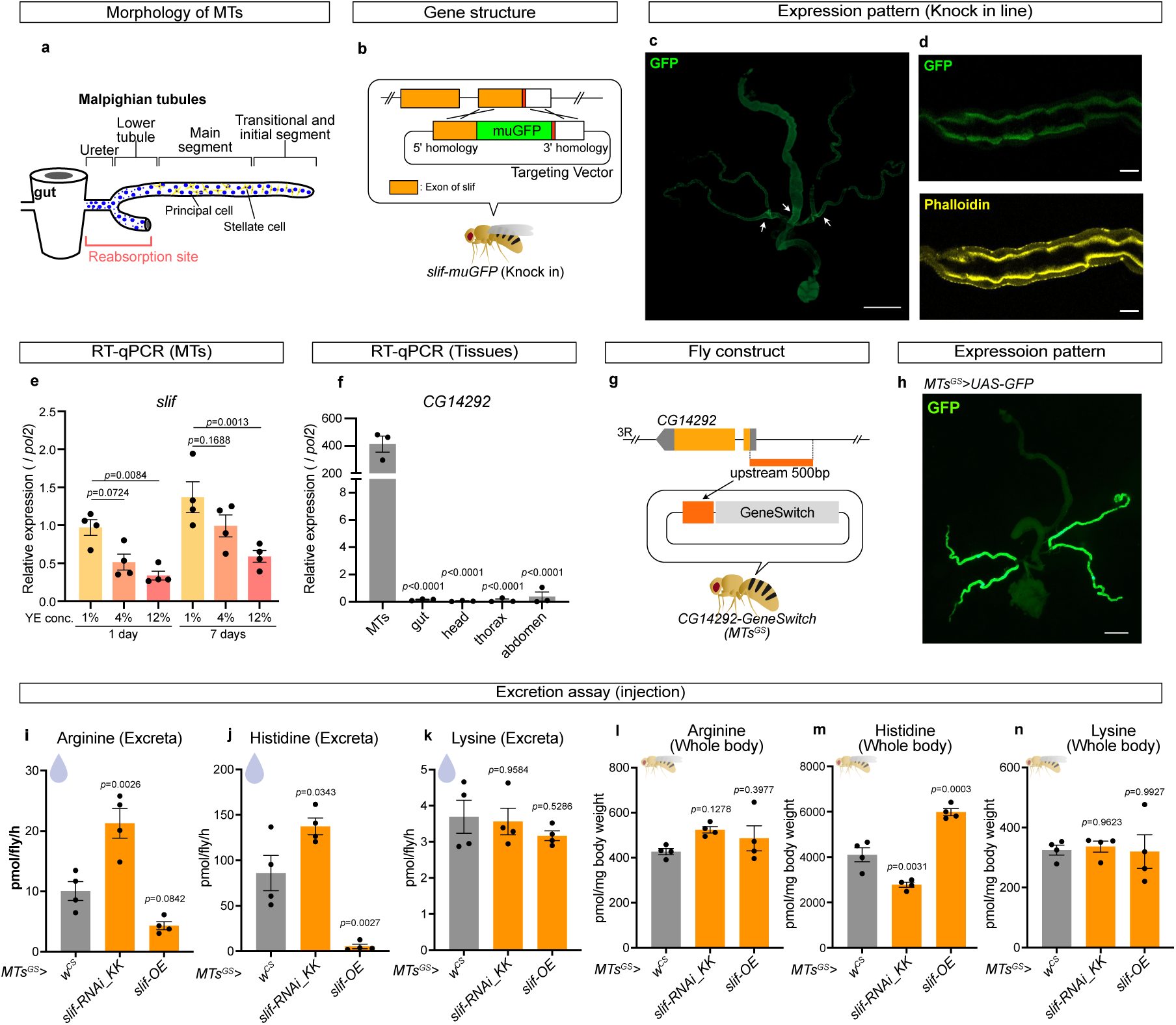
Slimfast-mediated amino acid reabsorption in the Malpighian tubules. **a** Schematic of MT regions. **b** Genetic structure of the *slif-muGFP* knock-in flies generated using the CRISPR-Cas9 system. Orange boxes, exons; white boxes, untranslated legions (UTRs); the green box, the muGFP sequence; and red boxes, stop codons. **c** Expression pattern of the Slif-muGFP in the gut and MTs. Scale bar: 500 µm. **d** Representative images of the cross-sectional view of the MTs (lower tubule). Green signals indicate muGFP (Slif) and yellow signals indicate phalloidin (actin). Scale bar: 20 μm. **e** RT-qPCR analysis of MTs from female CS flies fed diets containing YE at indicated concentration for 1 or 7 days. n = 4. Statistics: one-way ANOVA with Šidák’s multiple comparisons. **f** RT-qPCR analysis of tissues from 8-day-old female CS flies. n = 3. Statistics: one-way ANOVA with Šidák’s multiple comparisons. **g** Schematic of the generation of *MTs^GS^* (*CG14292^GS^*) flies. **h** GFP expression of *MTs^GS^>UAS-GFP* in the gut and MTs. Flies were fed a diet containing 50 μM RU486 for 3 days. Green, GFP signal. Scale bar: 500 µm. **i–n** Quantification of Arg, His, and Lys in excreta (**i–k**) and in whole bodies (**l–n**) of female flies of indicated genotypes fed a diet containing 200 μM RU486 for 6 days before the experiment. After injection with an AA cocktail, flies were placed in tubes for 4 h to collect excreta. n=4. Statistics: one-way ANOVA with Šidák’s multiple comparisons. Each graph shows the mean ± SEM.

To experimentally determine the spatial expression pattern of *slif* in the gut and MTs, we performed rolling circle amplification-based sequential fluorescent *in situ* hybridisation (sequential RCA-FISH). We examined the expressions of *Alp4*, *Drip*, *mex1*, *byn*, and *Mal-8A*, which are markers for the lower tubule, stellate cells, enterocytes, hindgut, and midgut R2 region, respectively. Corresponding signals for these genes were detected in the expected regions, validating the reliability of the method (Extended Data Fig. 9b). Using probes targeting *slif*, we detected the signals in the R5 region of the midgut and in the MTs, specifically in the ureter, lower tubule, and main segment, overlapping with regions marked by *Alp4* expression (Extended Data Fig. 9b). In females, the signals were also detected in the midgut (Extended Data Fig. 9b), possibly acting in the absorption of dietary AAs.

To examine subcellular localisation of Slif protein, we generated a knock-in fly line in which monomeric ultra-stable GFP (muGFP) was fused to the C-terminus of the endogenous *slif* sequence (Fig. 3b). The protein expression pattern was consistent with the RCA-FISH analysis; Slif-muGFP signal was strongly detected in the ureter and lower tubule regions (Fig. 3c). Higher magnification imaging of the lower tubule revealed that Slif localised to the apical membrane of tubule cells (Fig. 3d).

We next investigated whether *slif* expression responds to dietary AA levels. In our transcriptome analysis, *slif* transcription in MTs tended to increase upon His or Lys depletion, although the change was not statistically significant (Extended Data Fig. 8c). When we quantified *slif* expression by RT-qPCR analysis, we found that *slif* mRNA levels in MTs decreased as dietary YE concentration increased (Fig. 3e). Together, these results suggest that *slif* expression increases under low protein or AA deficient conditions and may promote the reabsorption of AAs.

### Slimfast is responsible for reabsorption of basic AAs

To investigate the function of Slif in the MTs, we generated a drug-inducible MTs-specific driver. Based on our transcriptome analysis, we identified *CG14292* as a gene highly and selectively expressed in the MTs and confirmed its enriched expression by RT-qPCR (Fig. 3f), although its function is unknown. We cloned a 500 bp upstream region of *CG14292* and created a fly line for GeneSwitch^34^ (Fig. 3g). When this driver line (*CG14292-GeneSwitch*, hereafter *MTs^GS^*) was crossed with *UAS-GFP*, strong GFP expression was induced in the MTs in an RU486-dependent manner (Fig. 3h and Extended Data Fig. 10a). Expression was observed throughout the MTs, including both principal and stellate cells (Fig. 3h and Extended Data Fig. 10b). Note that the expression of ureter was undetectable (Fig. 3h). Using this *MTs^GS^* driver, we manipulated *slif* and quantified AA excretion after injection of an AA cocktail. Strikingly, knockdown of *slif* increased Arg and His excretion and decreased internal His levels, while overexpression of *slif* produced the opposite effects (Fig. 3i, j, l, m). In contrast, Lys levels in the excreta and the bodies showed changed little under either condition (Fig. 3k, n). These results suggest that Slif transports only His and Arg, or that Slif may transport all three basic AAs but the effect on Lys levels is masked due to metabolic degradation of Lys in the MTs, as suggested in our *ex vivo* assay (Fig. 2g).Taken together, Slif appears to mediate the reabsorption of, at least, His and Arg from the tubule lumen back into the haemolymph.

### Slimfast transports basic amino acids

It has been suggested that Slif transports Arg^12^. However, there is no direct evidence that Slif can also transport His, nor how Arg/His transportation is regulated. To perform detailed transport analysis, we generated a stable *Drosophila* S2 cell line expressing *slif* under the control of a metallothionein (MT) promoter, which is inducible by Cu^2+^ (Fig. 4a). We confirmed that Slif localised to the plasma membrane (Fig. 4b) and was detected as a single band between 50 and 75 kDa by western blotting (Fig. 4c), consistent with its predicted molecular weight (66 kDa). Following Cu^2+^ induction, we examined Arg and His transport properties in Slif-expressing S2 cells using radioisotopes. Uptake of both [³H]L-Arg and [³H]L-His was significantly increased compared with non-induced controls and was inhibited by non-radiolabelled Arg or His, indicating that Slif transports both AAs (Fig. 4d, e). At pH 7.4, [³H]L-Arg transport did not require Na^+^, whereas [³H]L-His transport was increased in the presence of Na^+^ (Fig. 4d, e). [³H]L-Arg transport was unaffected by either Na^+^ or pH (Fig. 4f), indicating that Arg transport is Na^+^- and pH-independent. In contrast, His transport was increased markedly under acidic conditions (Fig. 4g, h). To test whether the Na⁺ dependence reflects a requirement for monovalent cations, we measured His transport at pH 7.0 in the presence of Na^+^ and K^+^. His transport increased in both Na^+^ and K^+^, demonstrating the importance of monovalent cations, not Na^+^ itself (Fig. 4i). Substrate selectivity was further examined using an inhibition assay of [³H]L-His transport in the presence of Na^+^ at pH 7.0, which confirmed that Slif preferentially transports cationic AAs over Ala, Leu, Phe, and Glu (Fig. 4j, k). Kinetic analysis revealed that His transport in the presence of Na⁺ at pH 7.0 exhibited a K_m_ of 115.8 μM and a V_max_ of 125.8 pmol/10^6^ cells/min (Fig. 4l). Furthermore, induction of *slif* expression by Cu²⁺ resulted in increased intracellular levels of the basic AAs Arg, His, and Lys (Fig. 4m), indicating that Slif broadly transports basic AAs, including Lys. Together, these findings identify Slif as a transporter capable of transporting basic AAs, whereas His transport exhibits pH sensitivity and is facilitated by monovalent cations at neutral pH.

**Fig. 4.**
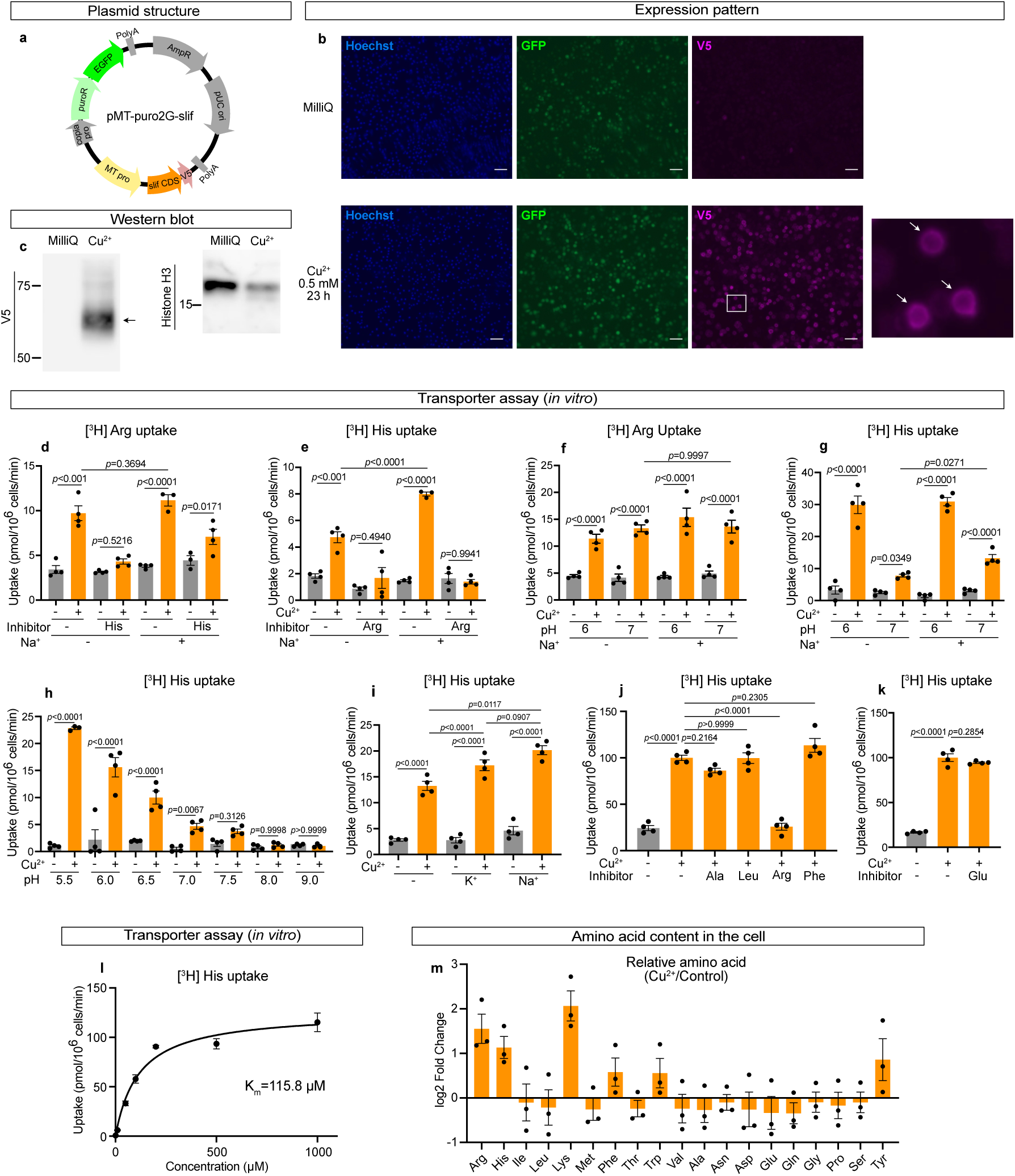
Functional characterisation of Slimfast in S2 cells. **a** Structure of the plasmid used to generate an S2 cell line stably expressing *slif*. **b** Representative image of a stable S2 cell line expressing *slif*. Blue signals indicate Hoechst 33342 (nuclei), green signals indicate GFP (puroR), and magenta signals indicate V5-tagged Slif. Scale bar, 50 µm. White arrows in the magnified image indicate the membrane localisation of Slif. **c** Western blot analysis of stable *slif*-expressing cells using an anti-V5 antibody. Anti-Histone H3 was used as a loading control. Arrow indicates the expected band for Slif. **d, e** Transport of [^3^H]L-Arg (**d**) or [^3^H]L-His (**e**) in the absence (-) or presence of the indicated non-radiolabelled AAs (3 mM) at pH 7.4. n = 3–4. **f, g** Transport of [^3^H]L-Arg (**f**) or [^3^H]L-His (**g**) in the absence or presence of Na^+^ at pH 6.0 or 7.0. n = 4. **h** pH dependence of [^3^H]L-His transport in the absence of Na^+^. n = 3–4. **i** Transport of [^3^H]L-His at pH 7.0 in the presence of different monovalent cations (K^+^ or Na^+^) using choline^+^ (-) as a control (n = 4). **j, k** Substrate selectivity of Slif determined by inhibition of [^3^H]L-His transport by the indicated AAs (Ala, Leu, Arg, and Phe (**j**) and Glu (**k**)) in the presence of Na^+^ at pH 7.0. n = 4. **l** Michaelis-Menten kinetics of [^3^H]L-His at substrate concentrations of 1-1,000 μM in the presence of Na^+^ at pH 7.0. n = 4. **m** Log2 fold changes in individual AA levels in stable *slif* expressing cells, normalised to the uninduced (control) group. For all experiments, MilliQ (control) or CuSO_4_ was supplemented to the medium over night before analysis. Each graph shows the mean ± SEM.

### Slimfast-dependent reabsorption is crucial for survival upon a high protein diet

Finally, we investigated the physiological significance of Slif by measuring survival curves in flies manipulating *slif* in the MTs. *slif* overexpression increased mortality under a high-protein diet containing 18% YE (Fig. 5a–d), and high-His diet containing 100 mM His (Fig. 5e–h). Control flies lacking the driver showed no survival difference with or without RU486 under either dietary condition (Fig. 5b, d, f, h), indicating that RU486 itself had no toxic effect. These results suggest that enhanced AA reabsorption increases susceptibility to high-AA stress. Overall, Slif-mediated AA reabsorption in MTs plays a pivotal role in maintaining organismal homeostasis in response to dietary AA levels (Fig. 5i).

**Fig. 5.**
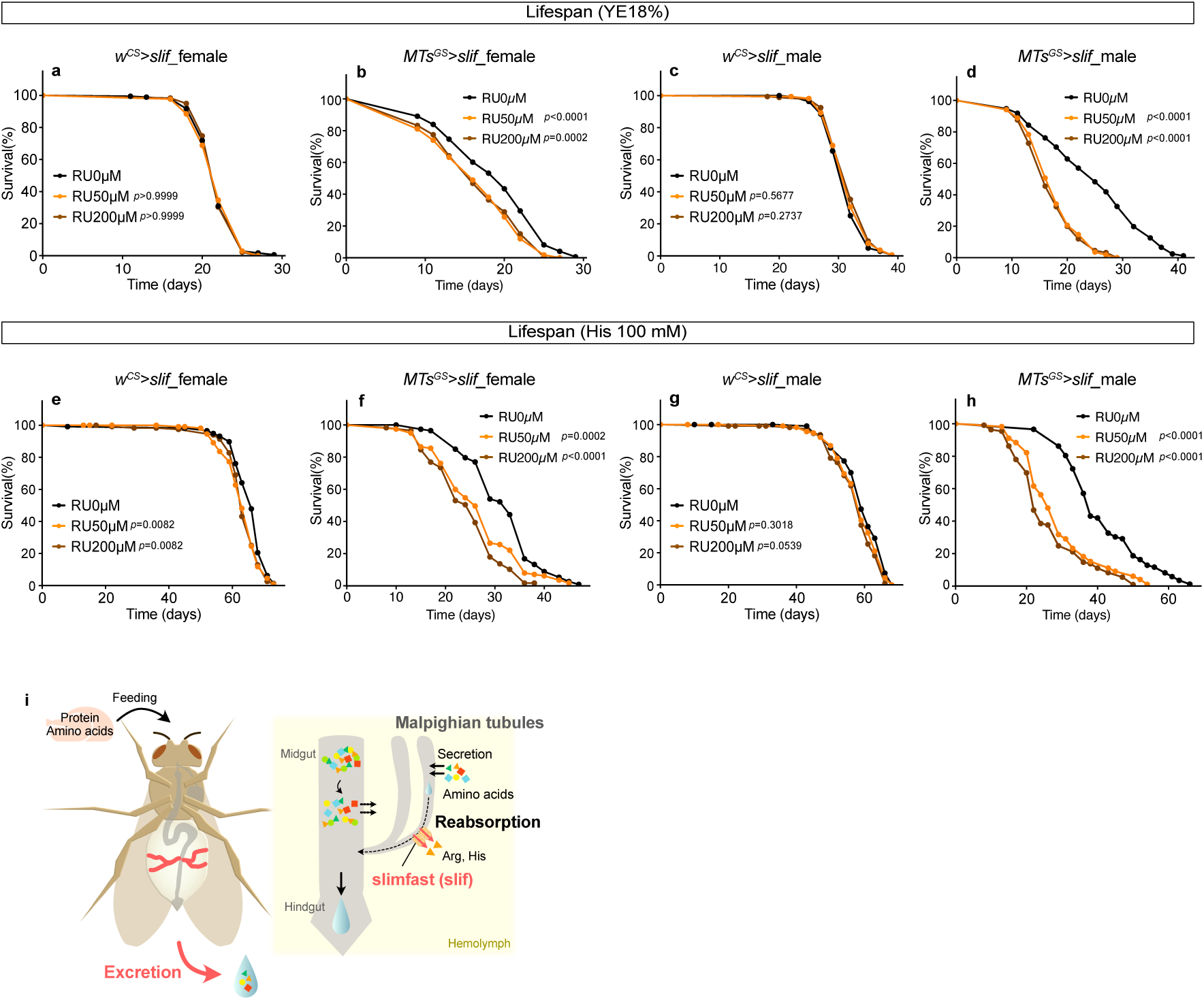
Amino acid reabsorption by Slimfast is crucial for survival under high protein diets. **a–d** Lifespan of female (**a, b**) and male (**c, d**) flies of indicated genotypes reared on 18% YE diet supplemented with indicated concentrations of RU486. n=181 for *w^CS^>slif*_RU 0 µM female, n=174 for *w^CS^>slif*_RU 50 µM female, n=178 for *w^CS^>slif*_RU 200 µM female, n=174 for *MTs^GS^>slif*_RU 0 µM female, n=169 for *MTs^GS^>slif*_RU 50 µM female, n=173 for *MTs^GS^>slif*_RU 200 µM female, n=166 for *w^CS^>slif*_RU 0 µM male, n=167 for *w^CS^>slif*_RU 50 µM male, n=173 for *w^CS^>slif*_RU 200 µM male, n=173 for *MTs^GS^>slif*_RU 0 µM male, n=171 for *MTs^GS^>slif*_RU 50 µM male, and n=177 for *MTs^GS^>slif*_RU 200 µM male. Statistics: log-rank test. **e–h** Lifespan of female (**e, f**) and male (**g, h**) flies of indicated genotypes reared on His 100 mM diet supplemented with indicated concentrations of RU486. n=117 for *w^CS^>slif*_RU 0 µM female, n=115 for *w^CS^>slif*_RU 50 µM female, n=118 for *w^CS^>slif*_RU 200 µM female, n=115 for *MTs^GS^>slif*_RU 0 µM female, n=119 for *MTs^GS^>slif*_RU 50 µM female, n=118 for *MTs^GS^>slif*_RU 200 µM female, n=113 for *w^CS^>slif*_RU 0 µM male, n=117 for *w^CS^>slif*_RU 50 µM male, and n=113 for *w^CS^>slif*_RU 200 µM male, n=87 for *MTs^GS^>slif*_RU 0 µM male, n=112 for *MTs^GS^>slif*_RU 50 µM male, and n=109 for *MTs^GS^>slif*_RU 200 µM male. Statistics: log-rank test. **i** The model of reabsorption-mediated regulation of AA homeostasis in *Drosophila* MTs.

## Discussion

In this study, we demonstrate in *Drosophila* that Slif in the MTs plays a crucial role in maintaining AA homeostasis by reabsorbing basic AAs into the haemolymph (Fig. 5i). Changes in AA excretion occur more rapidly and to a greater extent than changes in internal AA levels, indicating the tight regulation of AA excretion. Based on its localisation and function in MTs, we conclude that Slif mediates the reabsorption of basic AAs from the tubular lumen into the haemolymph. Enhanced AA reabsorption by overexpressing *slif* in MTs increased susceptibility to nutritional stress, indicating a crucial role of the transporter for metabolic homeostasis. These data suggest that AA reabsorption appears to be functionally important, even in *Drosophila* where glomerular filtration does not contribute to primary urine formation. Nitrogen derived from AAs is primarily excreted as uric acid in *Drosophila*^35^, which can form crystals in the tubular lumen under conditions such as inflammation or high-protein diets^36,37^. In this context, excretion of nitrogen in the form of soluble AAs prior to conversion into uric acid may represent an alternative strategy to avoid potential toxicity associated with crystal formation.

Our detailed analysis in S2 cells reveal that Slif transports Arg in a Na⁺- and pH-independent manner, whereas His transport depends on both Na⁺ and pH, with monovalent cations being particularly required under neutral pH conditions. The pH dependence of His transport is consistent with that of CAT1-mediated His transport reported upon its identification in mice^38^. Because CATs preferentially transport positively charged AAs, the properties of Arg and His transport can be explained by the distinct sidechain pKa values of these AAs. Arg (pKa ≈ 12.5) carries a +1 charge at pH 6–7.4 (experimental pH and physiological pH), whereas His (pKa ≈ 6.0) is largely neutral at pH 7.0–7.4 due to deprotonation of its imidazole sidechain and becomes predominantly positively charged under acidic conditions. Notably, the enhancement of His transport by monovalent cations at pH 7.0–7.4 suggests that these cations compensate for the lack of positive charge on His, thereby facilitating its transport under neutral conditions. His transport by Slif exhibited a K_m_ of 115.8 μM (Fig. 4l). Although the luminal concentration of His in the MTs is technically difficult to measure directly, our *ex vivo* assays indicate that approximately 40% of haemolymph His is excreted through the tubule lumen (Fig. 2g). Given that AA concentrations in larval haemolymph are in the mM range^39^, luminal His levels are estimated to fall within the hundreds to thousands of µM, exceeding the K_m_ of Slif. Based on these estimates, Slif operates near saturation under physiological conditions, such that His reabsorption is not limited by substrate availability. Instead, it is regulated by changes in transporter abundance, consistent with the observed diet-dependent regulation of *slif* expression (Fig. 3e), although the underlying molecular mechanism remains unclear. In the fat body, the membrane localisation of Slif is maintained under AA rich conditions through suppression of selective endocytosis downstream of mTORC1 signaling^40^. In mosquitoes, *Aa*Slif expression has been reported to be upregulated systemically following blood feeding, including in the fat body, gut, and MTs, suggesting a role in the efficient uptake of AAs derived from the blood meal^41^. While these studies collectively point to regulatory mechanisms that enhance AA uptake under nutrient rich conditions, our findings reveal that Slif may also promote AA reabsorption under AA-deprived conditions, suggesting the presence of alternative regulatory mechanisms.

There is growing evidence of mechanisms regulating AA excretion in mammalian kidneys. For example, disruption of the mTORC1 pathway in mouse renal tubules impairs endocytic capacity required for reabsorption, reduces transporter protein levels, and results in increased urinary excretion of glucose and AAs^42^. Proximal tubule specific deletion of ATF4 in mice alters the expression of multiple renal AA transporters and urinary and serum metabolite levels^43^. CAT1 (Slif ortholog) in mammalian cells is regulated by cytokines, growth factors, and hormones^44^. Under AA starvation, both CAT1 mRNA and protein levels are increased^45–47^, which is mediated by ATF4-dependent transcription^47,48^, as well as by the cap-independent translational control^49,50^. Although it remains unclear whether similar regulatory mechanisms operate in renal tubules *in vivo*, these findings indicate that transporter abundance and reabsorptive capacity can be dynamically modulated by systemic nutritional and metabolic cues. Of note, *CAT1* transcript was found in the renal distal convoluted tubules, the segment functions for fine-tuning ion composition in urine^51^. Given that subcellular localisation of CAT1 in polarised renal cells remains ambiguous (Human Protein Atlas; ^44,52^), our demonstration of Slif localisation and function in the MTs calls for revisiting the physiological role and regulatory mechanisms of CATs in mammalian renal system.

AAs regulate growth, reproduction, metabolism, behaviour and longevity in animals. In *Drosophila*, it has been widely studied how dietary intake, as well as metabolic regulation, of AAs impact various physiological responses to internal AA fluctuations^53,54^. Our present data show that excretory output, in addition to dietary input and internal metabolic conversion, forms an additional regulatory mechanism of AA homeostasis and provide a new strategy for improving healthspan. In human metabolic disorders such as phenylketonuria and maple syrup urine disease, mutations in metabolic enzymes lead to the accumulation of AA metabolites, some of which are not fully reabsorbed and are instead excreted in the urine^55,56^. In addition, defects in AA transporters in renal tubules have been reported to impair reabsorption, resulting in increased urinary AA levels and reduced systemic availability^57,58^. These observations suggest that nutrient reabsorption may be dynamically regulated to maintain homeostasis under different physiological conditions. Consistent with this concept, such regulatory principles are exemplified by the clinical use of SGLT2 inhibitors to modulate renal glucose reabsorption in treating diabetes. A deeper understanding of metabolite-dependent regulation of excretion may therefore provide new strategies for manipulating systemic nutrient balance. By utilising experimental advantages of the fly system, including lifespan analysis and precise manipulation of gene expression and diet, we can gain deeper insight into the role of excretion as a system for maintaining homeostasis in response to daily dietary fluctuations.

## Methods

### Fly stocks and husbandry

Flies were reared on a standard diet containing 4.5% cornmeal (NIPPN CORPORATION), 4% dry brewer’s yeast (Asahi Breweries, HB-P02), 6% glucose (Nihon Shokuhin Kako), and 0.8% agar (Ina Shokuhin Kogyo, S-6) with 0.4% propionic acid (Wako, 163-04726) and 0.03% butyl p-hydroxybenzoate (Wako, 028-03685). Flies were reared at 25 °C and 65% humidity with 12 h/12 h light/dark cycles. The fly lines used in this study were Canotn-S (CS), Dahomey, *w^Canton-S^* (*w^CS^*, *w^iso31^* eight times backcrossed with Canton-S), *UAS-2×eGFP* (Bloomington Drosophila Stock Center (BDSC), 6874), *UAS-GFP.nls* (BDSC, 4775), *UAS-slif-RNAi* (Vienna Drosophila Resource Center (VDRC), 110425), *UAS-slif* (BDSC, 52661).

Embryos were collected on an acetic acid agar plate (3% agar (Wako, 010-15815), 1% sucrose (Wako, 196-00015), and 0.3% acetic acid (Wako, 017-0256)) with live yeast paste (Oriental, 01402017) using a cage containing virgin female and male flies. Equal volumes of embryos were placed on the fly food. Eclosed flies were maintained for 2 or 3 days for maturation, and they were sorted by sex and put into vials (30 flies per vial). For manipulation using GeneSwitch, 50 or 200 μM RU486 (Tokyo Chemical Industry, M1732) dissolved in EtOH (Wako, 057-00451) or an equal volume of EtOH (as a control) was mixed in standard food.

### Dietary manipulations

For the YE diet, yeast extract (Nacalai Tesque, 15838-45) was mixed at the indicated concentrations with 6% glucose (Wako, 049-31165), 1% agar, 0.3% propionic acid, and 0.05% butyl p-hydroxybenzoate.

For the chemically defined diet (holidic medium), AA concentrations were adjusted according to the optimised diet through exome matching^25^. The composition of the holidic medium used in this study was described previously^21^. For diets restricted in His or Lys, all other ingredients, including AAs, were first mixed and aliquoted, and the restricted AA was added from pre-dissolved solutions.

### Generation of transgenic flies

To generate *slif-muGFP* knock in flies, we used CRISPR-Cas9 system^59^ to insert monomeric ultra-stable GFP (muGFP) at C terminus of the *slif* gene. An sgRNA target site of *slif* was selected using CRISPR Optimal Target Finder (http://targetfinder.flycrispr.neuro.brown.edu/)^60^. Complementary oligonucleotides with overhangs were annealed and cloned into the BbsI-digested U6b vector using a DNA ligation kit (Takara Bio, 6023). Sense strand: TTCGGACTACAAGGTCGAAAATGG and anti-sense strand: AAACCCATTTTCGACCTTGTAGTC. The targeting vector was constructed by inserting *pBluescript II SK (+)* with 500bp homology arms with a mutation in the PAM sequence next to the sgRNA site (from GGG to GAG). First, 500 bp homology arms were PCR-amplified from whole fly genome DNA by using PrimeSTAR^®^ Max DNA Polymerase (Takara Bio, R045). Primers for PCR were designed using the NEBuilder Assembly Tool (https://nebuilder.neb.com/#!/) and are listed in Table S2. The gel-purified PCR products were cloned into the EcoRI-digested *pBluescript II SK (+)* vector (Addgene, 212205) using NEBuilder HiFi DNA Assembly Master Mix (New England Biolabs, E2621X). The mixture of *pU6b-sgRNA* vector and targeting vector was microinjected into *w^1118^; attP40{nos-Cas9}/CyO* embryos by WellGenetics (New Taipei City, Taiwan). The F0 adults were crossed with balancer lines, and the muGFP-inserted lines were selected by genotyping. PCR primers for genotyping are listed in Table S2.

To generate *CG14292-GeneSwitch* (*MTs^GS^*) flies, the 500 bp sequence upstream of *CG14292* was PCR-amplified from whole fly genome DNA as a putative promoter and GeneSwitch sequence was PCR-amplified from *pelav-GeneSwitch* (Addgene, 83957) using PrimeSTAR Max DNA Polymerase. Primers for PCR were designed using the NEBuilder Assembly Tool and listed in Table S2. The gel-purified PCR products were cloned into the KpnI -digested *pelav-GeneSwitch* (Addgene, 83957) using NEBuilder HiFi DNA Assembly Master Mix. The vector was microinjected into *w^1118^* embryos by WellGenetics. F0 adults were crossed with balancer lines, and *w⁺*progenies were selected.

### Lifespan measurement

Thirty adult flies per vial were placed in a vial containing a fly diet. Dead flies were counted every 2 to 4 days when the flies were transferred to fresh vials.

### Fecundity analysis

Flies were quickly anaesthetised and separated into groups of three males with three females. After 24 hours incubation, the egg number in each vial was counted manually.

### Body and excreta collection in *in vivo* excretion assay

For the feeding assay, 1% Brilliant blue FCF (Wako Pure Chemical Industries, Ltd., 027-12842) was added to the experimental diet to visualise excretion, and flies were cultured for the indicated number of days. For the injection assay, the injection solution was first prepared. As stock solutions, we prepared an AA cocktail at 5× synthetic diet concentration, 10× PBS (Wako, 163-25265), and 10% Brilliant blue FCF. Each was mixed to achieve final concentrations of 4×, 1×, and 1%, respectively, and filled into glass capillaries prepared using a puller (NARISHIGE, PC-10). The capillary was attached to a Femtojet 4i (eppendorf), and the tip thickness was adjusted to produce droplets of 30-60 nl. The droplet was injected into the body cavity by inserting the capillary into the fly’s thorax. Four females and five males were placed in 1.5 ml tubes, covered with perforated Parafilm, and incubated at 25°C for the indicated duration.

For whole body samples, fly body weight was measured using a microbalance (METTLER TOLEDO, XR2UV), and the flies were homogenised in 160 µl of 80% methanol containing 10 µM internal standards (methionine sulfonate (Sigma, M0876-1G) and 2-morpholinoethanesulfonic acid (Dojindo, 341-01622)). For head samples, heads were carefully dissected and homogenised in 160 µl of 80% methanol buffer. For excreta samples, 300 µl of 80% methanol buffer was added to the tube and vortexed for 1 minute.

### Excreta collection in *ex vivo* excretion assay

A modified Ramsay assay^30^ was performed. Briefly, anterior MTs were carefully cut at the ureter in Schneider’s *Drosophila* medium (Thermo Fisher Scientific, 21720024). One side of the tubules was suspended in a 10-20 μl droplet of 1:1 *Drosophila* saline: Schneider’s *Drosophila* medium containing 0.2% Brilliant blue FCF (Wako, 027-12842), while the other side was wrapped around a metal pin. They were covered with fluid paraffin (Wako, 164-00476) to prevent evaporation. After 30-60 minutes of preincubation, excreta from the ureter were removed using a glass capillary, and the assay was started. After being left at room temperature for 2 hours, the excreta were imaged to measure their size, then aspirated using a pipettor fitted with a fine tip (Microloader) and collected into a tube containing 50 µl of 80% methanol buffer.

### Measurement of metabolites

Samples were centrifuged at 15000 ×g for 5 minutes, and the supernatant was collected. To remove proteins, 75 µl (for whole body and head), 150 µl (for excreta), and 25 µl (for ex vivo sample) acetonitrile (Wako, 015-08633) was added and centrifuged at 15000 ×g for 5 minutes. After passing through a Nanosep 10 kDa column (PALL, OD010C35) by centrifugation at 14000 ×g for 10 minutes, the samples were completely evaporated. They were dissolved in ultrapure water (Invitrogen, 10977-015) and injected into an LC-MS/MS (Shimadzu, LCMS-8060) with a PFPP column (Sigma, Discovery HS F5, 2.1 mm x 150 mm, 3 µm) in a column oven at 40 °C. A gradient from solvent A (0.1% formic acid in water) to solvent B (0.1% acetonitrile) was performed for 20 minutes. MRM methods for metabolite quantification were optimised using the software (Shimadzu, LabSolutions).

### RNA sequencing analysis for transcriptomics

Total RNA was extracted from 15-20 adult female MTs using a Promega ReliaPrepTM RNA Miniprep Kit (Promega, Z6112). Library preparation was performed by RIKEN BDR Genomics Research and Analysis Support Team using Illumina Stranded mRNA Prep, Ligation (96 Samples) (Illumina, 20040534) and IDT® for Illumina® RNA UD Indexes Set A/B Ligation (Illumina, 20040553/20040554). Optimal PCR cycle number was determined by real-time PCR using KAPA real-time library amplification kit (Roche, KK2702). The library quality was ensured by TapeStation HS D1000 assay. RNA sequencing was performed using an Illumina NovaSeq 6000. The paired-end 150 bp sequence data were analysed as follows: a quality check of the raw reads was performed by FastQC (v0.12.1). The raw reads were filtered to remove the adaptors and low-quality bases using Trim galore (v0.6.10). Filtered reads were aligned to the *Drosophila* genome (BDGP6.46) using Hisat2 (v2.2.1). The read counts were calculated using StringTie (v2.2.1). Differentially expressed genes were identified using DESeq2 (v1.44.0). Differentially expressed genes were defined as those with an adjusted P value (*p*_adj_) < 0.01 and a base mean expression > 50. Genes were further separated into upregulated (log2FC > 0) and downregulated (log2FC < 0) groups. Gene Ontology (GO) enrichment analysis was then performed separately for each gene set using the clusterProfiler package in R (function enrichGO) with the org.Dm.eg.db database. Over-representation analysis was conducted for Biological Process (BP) terms using FlyBase gene identifiers (keyType = “FLYBASE”). *P* values were adjusted using the Benjamini-Hochberg method and GO terms with adjusted *P* < 0.05 and q-values < 0.05 were considered significant. Enrichment results were visualised using dot plots generated with the clusterProfiler package. All RNA-seq results and GO enrichment results are provided in Table S1.

For heatmap, 64 putative AA transporter genes were extracted, and gene expression values were standardised by calculating z-scores across all samples for each gene. The z-score values were then averaged across biological replicates within each condition. Heatmaps were generated from the averaged z-scores using the pheatmap package in R, with hierarchical clustering applied to rows. Statistical significance was indicated within the heatmap.

### RT-qPCR analysis

Total RNA was extracted from 5-6 female MTs using Promega ReliaPrepTM RNA Miniprep Kit (Z6112). RT-qPCR was performed using a OneTaq RT-PCR kit (Promega, M0482S) with qTOWER3 (Analytik Jena). Pol2 was used as an internal control. Quantification was performed by the ΔΔct method. Primer sequences are listed in Table S2.

### Imaging analysis

MTs were dissected in PBS and fixed with 4% paraformaldehyde (PFA, Nacalai Tesque, 09154-14) for 30-60 minutes at room temperature. They were washed with PBST (PBS containing 0.1% Triton X-100 (Nacalai Tesque, 35501-15)) and incubated with 0.8 µM Hoechst 33342 (Invitrogen, H3570) and Phalloidin 647 (Abcam, 176759) for 30-60 minutes at room temperature. They were washed with PBST and mounted in SlowFade Gold (Invitrogen, S36936). Confocal images were obtained using a Leica SP8.

### Target sequence selection and padlock probe design for sequential RCA-FISH

Target RNA sequences for sequential RCA-FISH were selected using two complementary approaches: (i) a PaintSHOP (https://github.com/beliveau-lab/PaintSHOP_pipeline) pipeline-based design and (ii) a custom tiling-array-style design. Genome sequence, transcriptome sequence and gene models were obtained from FlyBase FB2024_01 (*D. melanogaster* Release 6.56), and analyses were restricted to chromosome sequences only. Five target sequences were selected per gene.

PaintSHOP is a Snakemake-based pipeline for genome-/transcriptome-scale oligonucleotide FISH probe design^61^. We ran PaintSHOP_pipeline v1.3 with the following parameters: ‘blockparse_min_length=26’, ‘blockparse_max_length=32’, ‘blockparse_min_tm=37’, ‘blockparse_max_tm=42’, and ‘model_temp=37’. From the isoform-resolved RNA probe sets, we retained only candidate sequences targeting constitutive exons shared by all isoforms of each target gene. To improve specificity, each candidate was screened against the complete *D. melanogaster* transcriptome sequences using BLAST+ (https://blast.ncbi.nlm.nih.gov/blast/Blast.cgi) ‘blastn’ (v2.12.0+) with ‘-task blastn-short’, and otherwise default parameters^62^. Candidates were retained only if the reverse-complement (target-binding) sequence produced no detectable BLAST hits to transcripts of genes other than the target.

For genes for which the number of PaintSHOP-derived target sequences did not reach five, additional target sequences were selected using a tiling-array-style approach. Because multiple isoforms can be transcribed from a single gene, the isoform showing the highest expression in MTs was selected as a representative isoform based on FlyAtlas2 data (https://motif.mvls.gla.ac.uk/FlyAtlas2/)^63^. This representative isoform sequence was then exhaustively tiled using a 30-nt sliding window with a 5-nt step size. Candidate oligonucleotides with melting temperatures (Tm) between 37 and 42 were retained, where Tm was computed identically to the PaintSHOP pipeline implementation (Biopython ‘Tm_NN’ followed by ‘chem_correction’ with the same parameterisation). Candidate sequences were screened against the full *D. melanogaster* transcriptome sequences using the same BLAST+ procedure as described above, and only sequences without detectable off-target hits were retained.

The PaintSHOP-derived and tiling-derived candidate sets were merged for each gene. The additional tiling-derived candidates were selected such that the distances between target sequences along the gene were evenly distributed. For genes still failing to reach the required number of target sequences after combining additional tiling-derived candidates were selected by prioritising those with the lowest BLAST similarity to other genes.

Selected target sequences were converted into padlock probes (PLPs) by splitting each target sequence into two adjacent target-complementary arms, separated by a gene-specific ID sequence and a universal anchor sequence. Five PLPs targeting the same gene shared an identical gene-specific ID sequence. The universal anchor sequence was shared among all PLPs and was adopted from the HybISS protocol^64^.

Gene-specific ID sequences (GSIDs) were designed *in silico* as follows. Sequence filtering thresholds were matched to those used in the HybISS protocol^64^, except for the homopolymer criterion. A pool of five million fully random 20-nt sequences was generated and filtered to remove candidates containing homopolymer runs of ≥4 identical nucleotides, to enforce a local GC fraction of 40–50% within any 10-nt sliding window, and to retain candidates with Tm between 47.0 and 50.5 ℃. The resulting set was treated as the raw GSID candidate library. To ensure sequence orthogonality within the bridge probe library, each GSID candidate was screened against the raw GSID candidate library using BLAST+ blastn with ‘-task blastn-short’, using default parameters otherwise. Candidates were retained only if no hits were detected to other GSID candidates, or if the longest alignment to any other GSID candidates was <10 nt. To minimize secondary-structure formation, minimum free energy (MFE) was computed for each sequence using RNAfold v2.6.4 (https://www.tbi.univie.ac.at/RNA/)^65^, and candidates with MFE ≥ −7 kcal/mol were retained. Finally, 3,661 GSIDs remained, and six of these were randomly selected for use in the experiment.

The PLPs were synthesized as pooled oligonucleotides (oPools Oligo Pools, Integrated DNA Technologies) and 5′-phosphorylated using T4 polynucleotide kinase (Takara, 2021A) before use. Bridge and detection probe sequences were designed based on the HybISS protocol^64^. Three detection probes conjugated with Alexa Fluor 488, Alexa Fluor 555, or Alexa Fluor 647 at the 5′ terminus were synthesized by (Thermo Fisher Scientific). All probe sequences were listed in Table S3. In addition to the 30 PLPs targeting the six genes analysed in this study, the pooled PLP mixture included 405 additional PLPs targeting the 81 other genes. However, signals from these genes were not imaged.

### mRNA visualisation by Sequential RCA-FISH

The Sequential RCA-FISH was performed based on the RCA-based *in situ* sequencing protocol^64,66,67^, with modifications. In this study, combinatorial barcoding was not used for gene identification. Instead, in each imaging round, three bridge probes were used, each corresponding to a distinct gene–detection probe pair, such that each gene was assigned to a specific fluorescence channel. Thus, three genes were detected per imaging round.

Adult foregut, midgut, hindgut and MTs were dissected in PBS and mounted onto glass-bottom dishes (Matsunami glass, D11130H). Samples were then fixed with 4% PFA for 20 min at room temperature, followed by washing with PBSwx (1× PBS, 0.2 % TritonX-100, and 0.2% Tween-20). Samples were subsequently equilibrated in a 1:1 mixture of PBSwx and pre-hybridisation buffer (2× SSC, 20% formamide, 50 µg/mL heparin sodium salt, 0.2 % TritonX-100, 0.2% Tween-20 and 20 mM Ribonucleoside Vanadyl Complex (New England Biolabs, S1402S)) at 37 °C for 5 min, followed by incubation in pre-hybridisation buffer at 37 °C for 10 min. PLP hybridisation was then carried out in PLP hybridisation mixture (2× SSC, 20% formamide, 20 mM Ribonucleoside Vanadyl Complex, 50 µg/mL heparin sodium salt, 0.2 mg/mL BSA, 0.2 % TritonX-100, 0.2% Tween-20, and pooled PLPs at 0.01 µM per oligo) at 37 °C for 17 hours. Samples were washed with post-hybridisation buffer (2× SSC, 20% formamide, 0.2 % TritonX-100, 0.2% Tween-20 and 20 mM Ribonucleoside Vanadyl Complex) at 30 °C for 30 min, twice. After washing with PBSwx, PLP ligation was performed using the PLP ligation mixture (1× SplintR ligase reaction buffer, 0.2% Triton X-100, 0.2% Tween-20, 0.2 U/µL SUPERase·In RNase Inhibitor (Invitrogen, AM2696), and 1 U/µL SplintR Ligase (New England Biolabs, M0375L)) at 25 °C for 3 h. After washing with PBSwx, primer hybridisation was performed in primer hybridisation mixture (1× PBS, 50 µg/mL heparin sodium salt, 0.2 % TritonX-100, 0.2% Tween-20, 20 mM Ribonucleoside Vanadyl Complex, and RCA primer at 0.1 µM) at room temperature for 1 hour, followed by washing with PBSwx. RCA was performed in RCA mixture (1× Phi29 DNAP reaction buffer, 5% glycerol, 0.25 mM dNTP, 0.2 mg/mL BSA, 0.2% Triton X-100, 0.2% Tween-20, 0.2 U/µL SUPERase·In RNase Inhibitor, and 1 U/µL Phi29 DNA polymerase (Monserate, 4002)) at 30 °C for 17 hours, followed by washing with PBSwx. Then, samples were post-fixed in 4% formaldehyde (Sigma-Aldrich, 252549) in 1× PBS at room temperature for 10 min and washed with PBSwx. Samples were subsequently equilibrated in a 1:1 mixture of pre-hybridisation buffer and PBSwx at room temperature for 5 min, followed by incubation in pre-hybridisation buffer alone at room temperature for 10 min.

To labell the RCA products of three genes per imaging round with fluorescent probes, three bridge probes were hybridised in bridge probe hybridisation mixture (2× SSC, 20% formamide, 50 µg/mL heparin sodium salt, 0.2 % TritonX-100, 0.2% Tween-20, and three bridge probes at 0.1 µM each) at room temperature for 1 hour. Then, samples were washed with 2× SSCwx (2× SSC, 0.2% Triton X-100, 0.2% Tween-20) and subsequently incubated in detection probe mixture (2× SSC, 20% formamide, 50 µg/mL heparin sodium salt, 0.2% Triton X-100, 0.2% Tween-20, and three detection probes at 0.1 µM each) at room temperature for 1 h, followed by washing with 2× SSCwx. After washing with PBSwx, Nuclei were stained with DAPI solution (1× PBS, 0.2% Triton X-100, 0.2% Tween-20, and 1 µg/mL DAPI (DOJINDO, 340-07971)) at room temperature for 30 min and washed with PBSwx. Samples were imaged in PBS using a spinning-disk confocal microscope (Olympus IX81-ZDC2 equipped with a Yokogawa CSU-W1 spinning-disk unit and a Hamamatsu Photonics ORCA-Fusion BT camera) with a 20× air objective (Olympus UPlanXApo 20×, NA 0.8). Lasers at 405, 488, 561, and 650 nm were used for excitation. In addition to the four fluorescence channels, transmitted-light images were also acquired. To cover the entire foregut, midgut, hindgut, and MTs, tiled imaging was performed using a motorised stage and MetaMorph software (Ver. 7.10.5.476), with 10% overlap between adjacent tiles. For each tile, z-stack images were acquired at 1 µm intervals across 101 optical sections.

In this study, *mex1*, *Drip*, and *byn* transcripts were fluorescently labelled and imaged in the first imaging round. After imaging, samples were washed with 2× SSCwx and incubated in stripping buffer (2× SSC, 65% formamide, 0.2% Tween-20, 0.2% Triton X-100) for 1 h at room temperature, with the buffer replaced after 30 min. After washing with 2× SSCwx, the second imaging round was performed to detect *Alp4*, *Mal-A8*, and *slif* transcripts as described above.

### Sequential RCA-FISH image analysis

For each imaging round, tile, and channel, maximum-intensity projection (MIP) images were generated from the corresponding z-stacks using custom Python scripts. The resulting five-channel MIP images (DAPI, Alexa Fluor 488, Alexa Fluor 555, Alexa Fluor 647, and transmitted-light) were then merged into multi-channel images on a per-tile basis. For each imaging round, tiled images were stitched in Fiji/ImageJ (ImageJ 1.54p) using the Grid/Collection stitching plugin (Ver. 3.1.9, https://imagej.net/plugins/grid-collection-stitching), with default parameters. Stitched images from sequential imaging rounds were subsequently aligned by inter-round registration in Fiji using the Descriptor-based registration (2D/3D) plugin (Ver. 2.1.8, https://imagej.net/plugins/descriptor-based-registration-2d-3d) with a rigid (2D) transformation model, using the DAPI channel as the reference. The round 2 DAPI image was used as the fixed reference, and the estimated transformation was applied to all channels of round 1to generate registered multi-channel images. Brightness and contrast were adjusted individually for each sample and gene for visualization purposes, while ensuring that fluorescence signals were not saturated.

### Generation of stable *slif-*expressing cell lines

*Drosophila* S2 cells were maintained at 25°C in Schneider’s *Drosophila* medium supplemented with 10% (v/v) heat-inactivated fetal bovine serum (Thermo Fisher Scientific, SH30910.03) and 1×Penicillin-Streptomycin-L-Glutamine Solution (Wako, 161-23201). To generate vectors containing *slif* sequence, the coding sequence of *slif* was amplified by PCR using MTs cDNA as a template. The backbone vectors *pMT-PURO2G* (RDB09004) and *pMT-PURO2R* (RDB09005) were provided by the RIKEN BRC through the National BioResource Project of the MEXT/AMED, Japan. They were linearised by PCR using primers (listed in Table S2) designed by the NEBuilder Assembly Tool. The PCR-amplified *slif* insert and the linearised backbone vectors were assembled using the NEBuilder HiFi DNA Assembly Master Mix. Cells were transfected with each vector using Effectene Transfection Reagent (QIAGEN, 301425) according to manufacturer’s instructions. Briefly, 1×10^6^ cells were seeded in 6-well plates and treated with 0.4 μg DNA. After 24 hours of transfection, medium was replaced to 600 μl Schneider’s *Drosophila* medium without serum and incubated for another 24 hours. To select the transfected cells, medium containing 2 µg/ml puromycin (Wako, 166-23153) was used. Selection was continued for more than 6 weeks and slif was expressed in over 80% of cells upon induction of the MT promoter by 0.5 mM CuSO_4_ (Wako, 033-04415).

### Immunocytochemistry

Stable slif-expressing S2 cells were seeded in 6-well plates and treated with 0.5 mM CuSO₄ overnight at 25°C to induce expression. Cells were then plated onto coverslips coated with concanavalin A (0.5 mg/mL, Sigma, C5725) and incubated for 2 hours at 25°C. Cells were fixed with 4% PFA for 30 minutes at room temperature, followed by washing with PBST (PBS containing 0.1% Triton X-100 (Nacalai Tesque, 35501-15). After blocking with PBST containing 5% normal donkey serum, cells were incubated with anti-V5 antibody (1:200, Sigma, R960-25, RRID: AB_255656) diluted in PBST for 1 hour at room temperature. Samples were washed with PBST and incubated for 1 hour at room temperature with donkey anti-mouse IgG (H+L) highly cross-adsorbed secondary antibody conjugated to Alexa Fluor Plus 488 (1:500, Invitrogen, A31570, RRID: AB_2536180) and 0.8 μM Hoechst 33342 in PBST. Finally, samples were mounted in SlowFade Gold, and images were acquired using an Axio Vert.A1 microscope (ZEISS Microscopy, Jena, Germany).

### Cell collection for metabolite measurement

Stable slif-expressing S2 cells were treated with 0.5 mM CuSO₄ and incubated overnight at 25°C. After washing with PBS, cells were harvested in 160 μL of 80% methanol buffer and subjected to metabolite extraction followed by LC-MS/MS analysis as described above.

### *In vitro* transporter assay

AA transport was measured in stable *slif*-expressing S2 cells, which were maintained in 6 µg/mL puromycin. One day before the analysis, *slif* expression was induced by 0.5 mM CuSO_4_ and the cells without Cu^2+^ induction were used as negative controls. For each transport reaction, 1×10^6^ cells were used. The cells were washed twice with the assay buffer (Na^+^-free Hanks’ balanced salt solution (HBSS): 25 mM HEPES, 125 mM choline-Cl, 4.8 mM KCl, 1.2 mM MgSO_4_, 1.2 mM KH_2_PO_4_, 1.3 mM CaCl_2_, 5.6 mM D-glucose; or indicated buffers) and the medium was replaced with the assay buffers. Transport of 10 µM [^3^H]L-Arg (3.7 GBq/mmol) or 10 µM [^3^H]L-His (1.85 GBq/mmol) was measured for 2 min at 25 °C in the assay buffers (Na^+^-free HBSS or Na^+^-containing HBSS (choline-Cl replaced with NaCl)) containing radioisotope substrates at the indicated pH. For inhibition experiments, test amino acids were added together with the radioisotope substrates. After 2-min incubation, the reactions were terminated by the addition of ice-cold assay buffers, and cells were collected on GF/B glass microfiber filtration (Cytiva), followed by washing once. The filters were mixed with Emulsifier-Safe scintillation cocktail (Revvity) and radioactivity was measured by a β-scintillation counter (Tri-Carb 3110TR; PerkinElmer).

For pH dependent experiment, MES (for pH 5.5, 6.0, and 6.5) or Tris (for pH 8.0 and 9.0) were used instead of HEPES. In the cation selectivity experiments, choline-Cl was substituted with KCl or NaCl as indicated. The [^3^H]L-His transport kinetics were determined for 2 min at concentrations of 1, 10, 50, 100, 200, 500, and 1,000 μM at pH 7.0 in the presence of Na^+^. The transport values were fitted to the Michaelis-Menten equation using nonlinear regression.

### Western blot analysis

Stable slif-expressing S2 cells were seeded in 6-well plates and treated with 0.5 mM CuSO₄ overnight at 25°C to induce expression. The culture medium was aspirated, and cells were directly lysed on the plate with 200 μL RIPA buffer (Wako, 188-02453) supplemented with protease inhibitor (Wako, 165-26021). Lysates were clarified by centrifugation (15,000 × g for 5 min at 4°C), and protein concentrations were determined using a BCA assay (Wako, 164-25935). The samples were mixed with 6× SDS-PAGE sample buffer (Nacalai Tesque, 09499-14) and boiled at 98°C for 5minutes. 5 μg protein samples were subjected to standard SDS-PAGE using 10-20% Gel (Wako, 198-15041). Gels were transferred to the PVDF membrane and blocked with EveryBlot Blocking Buffer (Bio-Rad, 12010020). Primary antibodies used in the study were mouse anti-Histone H3 (1:1000, Cell Signaling Technology, 14269, RRID: AB_2756816) and mouse anti-V5 (1:1000). For the secondary antibody, Horseradish peroxidase (HRP)-conjugated secondary antibodies, anti-mouse IgG, HRP-linked antibody (1:1000, Cell Signaling Technology, 7076S, RRID: AB_330924) was used. The signals were visualised by chemiluminescence using Immobilon (Millipore, WBLUF0100) and detected using an Amersham ImageQuant 800 imager (Cytiva, Tokyo, Japan).

### Statistical analysis

Statistical analysis was performed using GraphPad Prism 10, 11 and R. A two-tailed Student’s t test was used to test between two groups. One-way ANOVA with Sidak’s multiple comparisons was used to compare multiple groups. The log-rank test was used for survival analysis. Log rank test with holm correction was used for multiple comparisons. All the statistical details can be found in the figure legends. All experimental results were repeated at least twice to confirm the reproducibility. Bar graphs were drawn as the mean and SEM with all the data points shown by dots to allow readers to see the number of samples and each raw data point.

## Acknowledgements

We thank the Kyoto Stock Center, National Institute of Genetics (NIG), Vienna Drosophila Resource Center (VDRC) and Bloomington Drosophila Stock Center (BDSC) for *Drosophila* stocks; RIKEN BioResource Research Center (BRC) for plasmid stocks; RIKEN BDR Genomics Research and Analysis Support Team for supporting RNA-seq analysis; Dr. Tadashi Uemura for critical comments on the manuscript; Mr. Housei Wada for supporting image analysis of RCA-FISH; Dr. Koshi Imami and Ms. Kaho Takamuro for generating data related to this study that are not included in this manuscript; Dr. Naoki Okamoto for sharing a transporter list and for critical advice; all members of Miura’s lab and Obata’s lab for technical assistance and important advice. This work was supported by The Japan Society for the Promotion of Science to F.O. (23K24032 and 26H02320), A.O. (25K23787 and 26K18471), P.W. (25K01961), S.N. (24K02888), and T.K. (22H05167), AMED-PRIME to F.O. (20gm6310011), JST FOREST Program to F.O. (JPMJFR2337), The Uehara Memorial Foundation to A.O., and RIKEN Special Postdoctoral Researchers program to A.O.

## Author contributions

F.O. and A.O. conceived the project; A.O. performed most of the experiments and analysed the data; S.N., P.W., and H.H. performed *in vitro* transporter assays; A.M., M.K., Y.K, and C.S. supported the experiments; T.K., D.C. and O.N. performed sequential RCA-FISH analysis; A.O. and F.O. wrote the initial manuscript; M.M. and F.O. supervised the study; All authors edited and approved the final manuscript.

## Data availability

RNA-sequencing data have been deposited at DDBJ under accession number PRJDB40568. All the materials generated in this study are available upon request to F.O.

---

**Extended Data Fig. 1.**
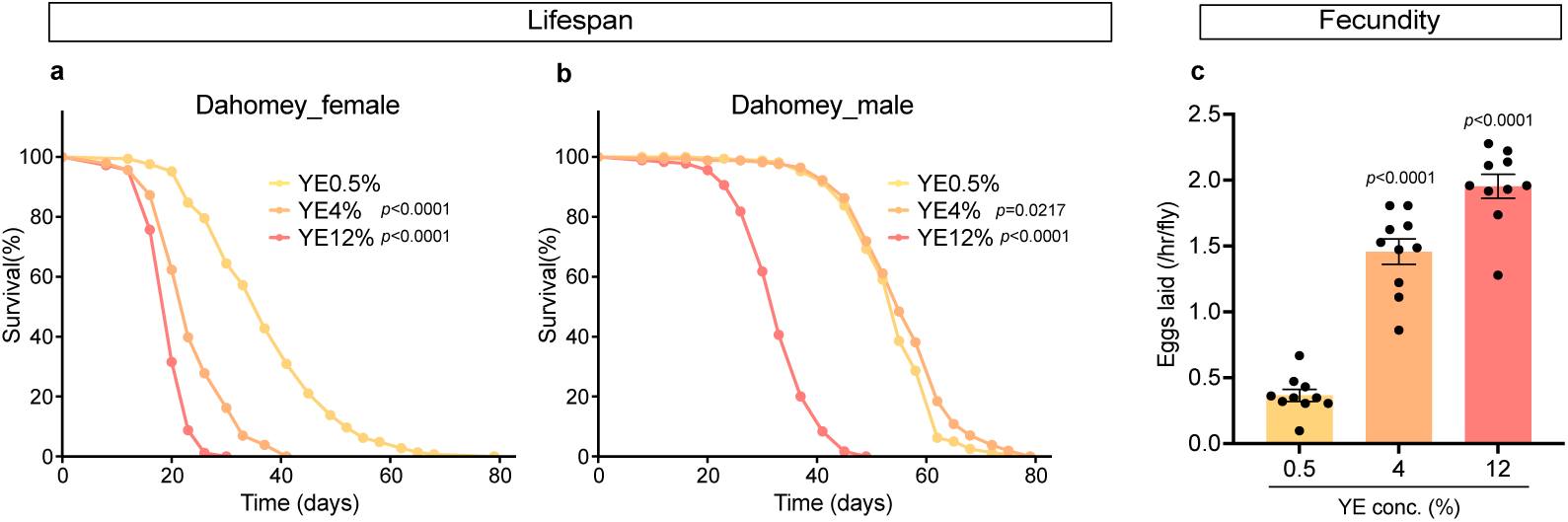
Diet-dependent response in Dahomey strain. **a, b** Lifespan of female (**a**) and male (**b**) Dahomey flies reared on diet containing yeast extract (YE) at indicated concentrations. n=167 for YE 0.5% female, n=189 for YE 4% female, n=183 for YE 12% female, n=173 for YE 0.5% male, n=176 for YE 4% male, and n=182 for YE 12% male. Statistics: log-rank test. **c** Fecundity of female Dahomey flies fed a diet containing YE at indicated concentrations for 1 day. n=10. Statistics: one-way ANOVA with Šidák’s multiple comparisons. Each graph shows the mean ± SEM.

**Extended Data Fig. 2.**
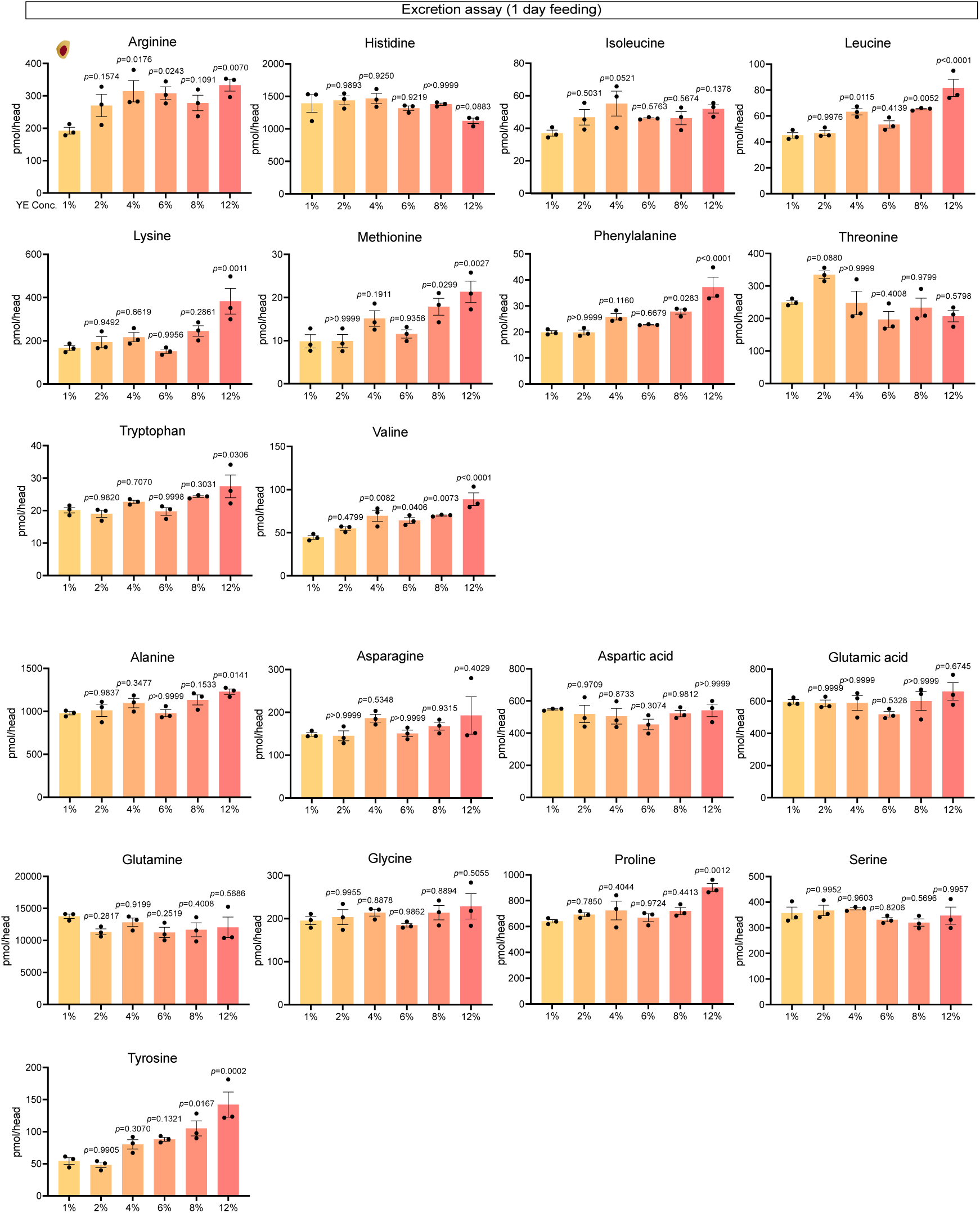
Internal levels of individual amino acids 1 day after YE feeding. Quantification of individual AAs in the heads of female CS flies after feeding on diets containing the indicated concentrations of YE for 1 day, followed by 4 h incubation in tubes. n=3. Statistics: one-way ANOVA with Šidák’s multiple comparisons. Each graph shows the mean ± SEM.

**Extended Data Fig. 3.**
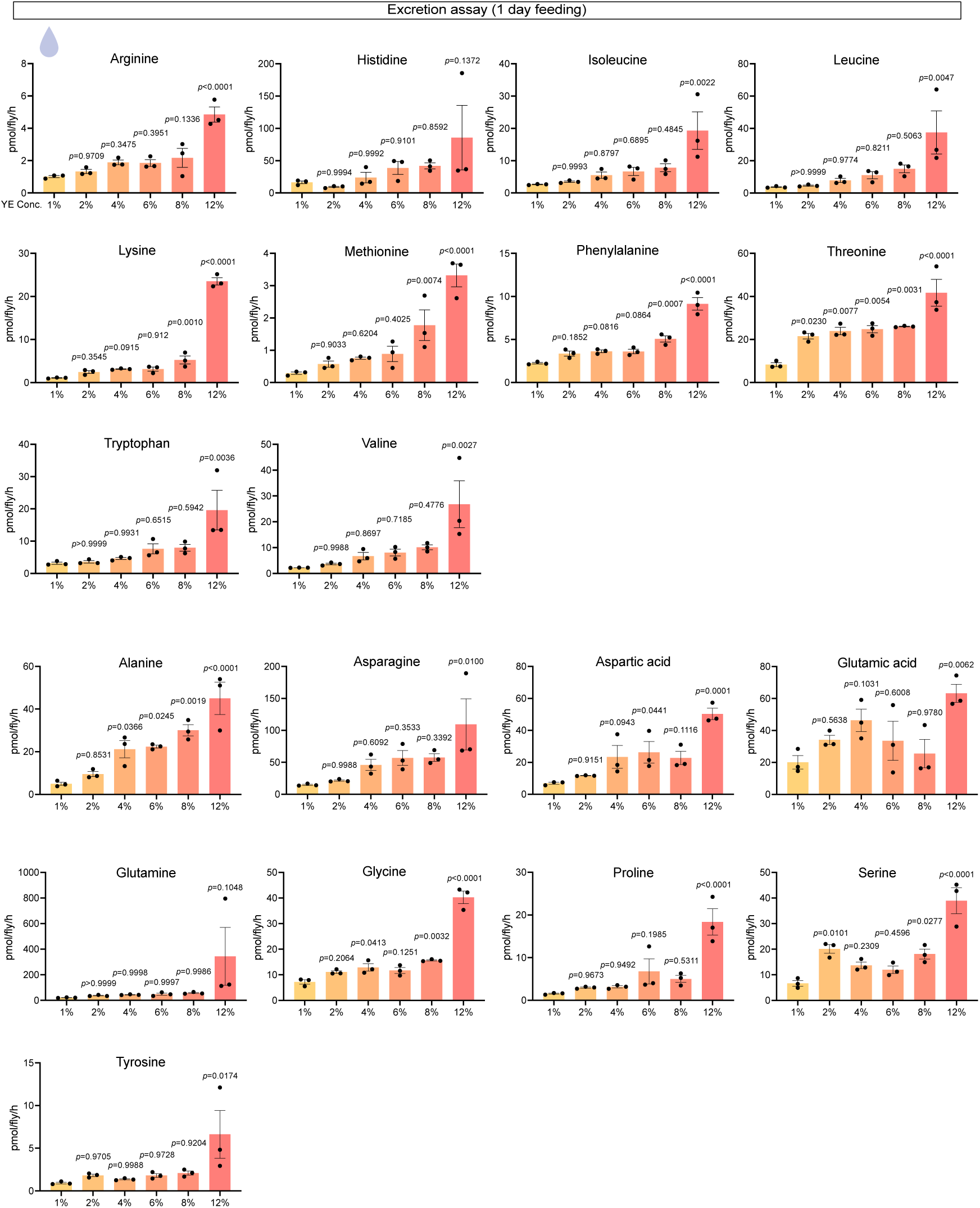
Excretion levels of individual amino acids 1 day after YE feeding. Quantification of individual AAs in the excreta of female CS flies after feeding on diets containing the indicated concentrations of YE for 1 day, followed by 4 h incubation in tubes. n=3. Statistics: one-way ANOVA with Šidák’s multiple comparisons. Each graph shows the mean ± SEM.

**Extended Data Fig. 4.**
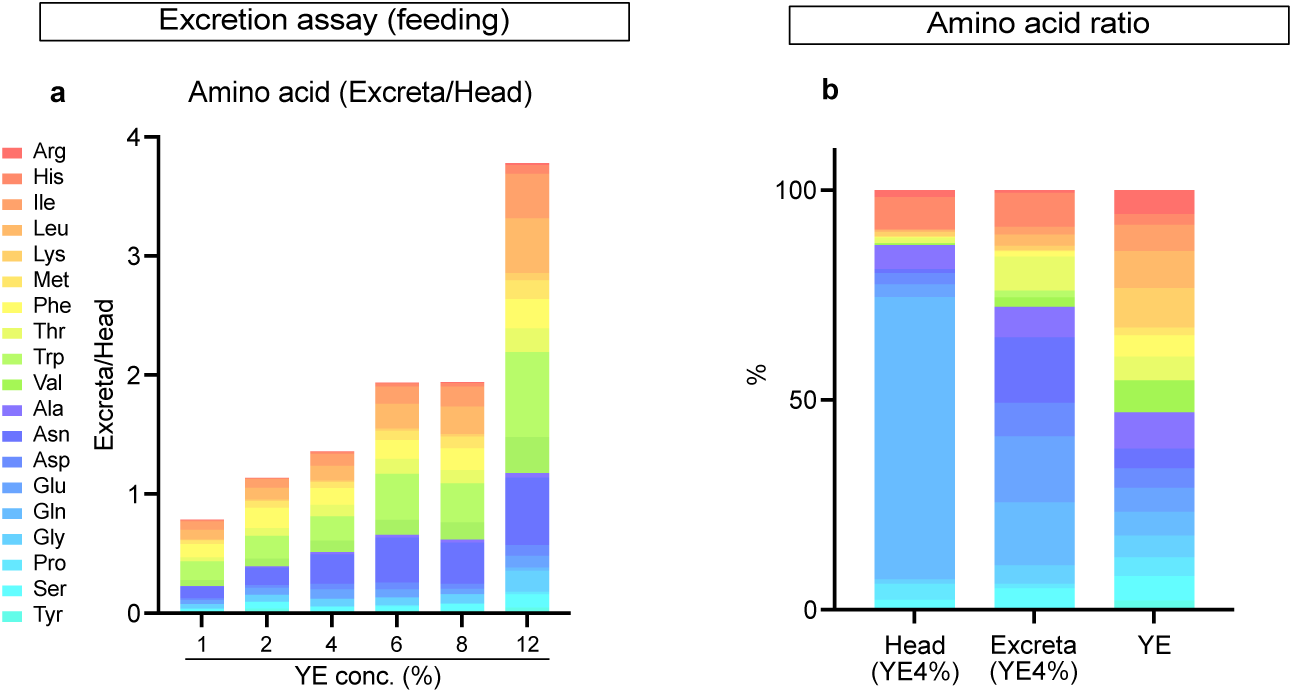
Calculated amino acid ratio in bodies and excreta. **a** Ratio of AA in excreta relative to head AA levels. **b** The AA composition of head, excreta, and YE diet, respectively.

**Extended Data Fig. 5.**
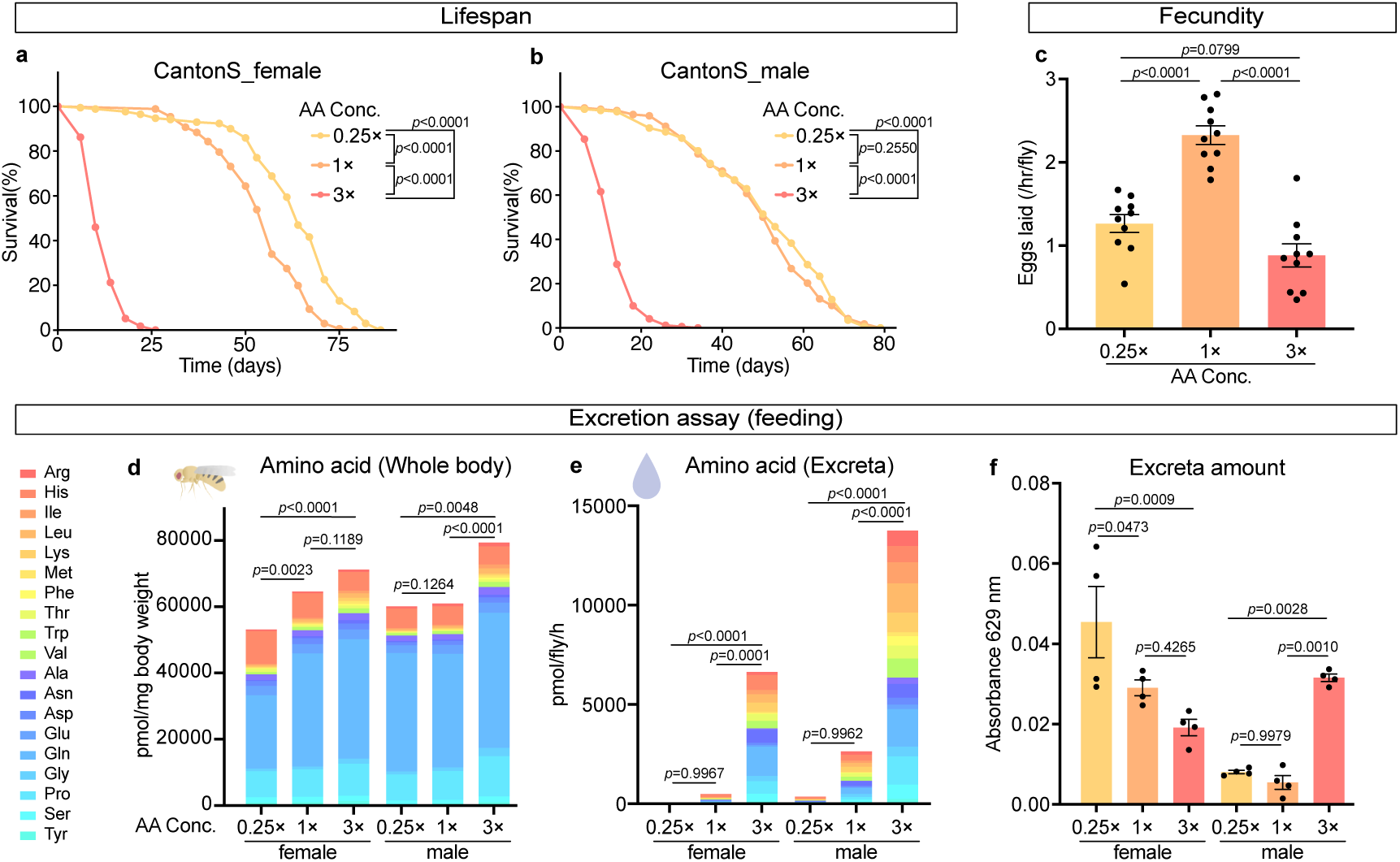
Lifespan, fecundity, and excretion in response to dietary amino acid levels. **a, b** Lifespan of female (**a**) and male (**b**) CS flies reared on synthetic diets containing AAs at indicated concentrations. n=170 for AA 0.25× female, n=173 for AA 1× female, n=174 for AA 3× female, n=176 for AA 0.25× male, n=170 for AA 1× male, and n=177 for AA 3× male. Statistics: log-rank test. **c** Fecundity of female CS flies reared on synthetic diets containing AAs at indicated concentrations for 1 day. n=10. Statistics: one-way ANOVA with Šidák’s multiple comparisons. **d, e** Quantification of AAs in whole bodies (**d**) and excreta (**e**) in female and male CS flies after feeding on synthetic diets containing AAs at indicated concentrations, followed by 2 h incubation in tubes. n=4. Statistical analysis was performed using the total AA levels. Statistics: one-way ANOVA with Šidák’s multiple comparisons. **f** Absorbance at 629 nm of excreta remaining in the tubes and solubilised in buffer. n=4. Statistics: one-way ANOVA with Šidák’s multiple comparisons. Each graph shows the mean ± SEM.

**Extended Data Fig. 6.**
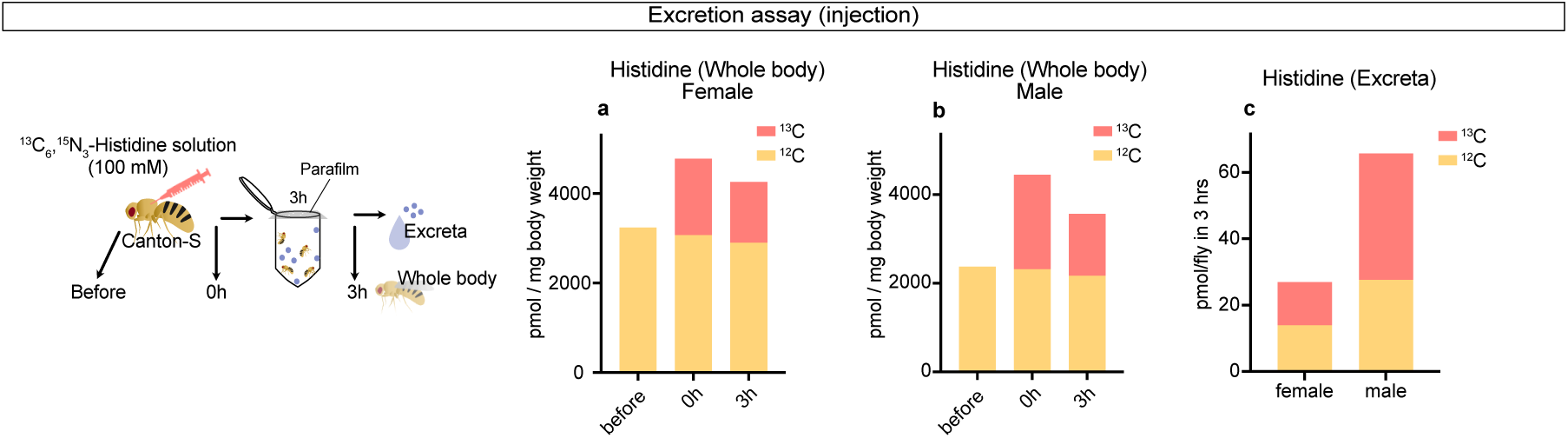
Injected amino acids are excreted from the haemolymph. **a, b** Quantification of ^12^C and ^13^C His in whole bodies of female (**a**) and male (**b**) CS flies before and after injection with^13^C His solution. n=2 for before and 0 h, and n=3 for 3 h after injection. **c** Quantification of ^12^C and ^13^C His in excreta of male and female CS flies 3 h after injection with ^13^C His solution. n=3. Statistics: unpaired two-tailed Student’s t test. Each graph shows the mean ± SEM.

**Extended Data Fig. 7.**
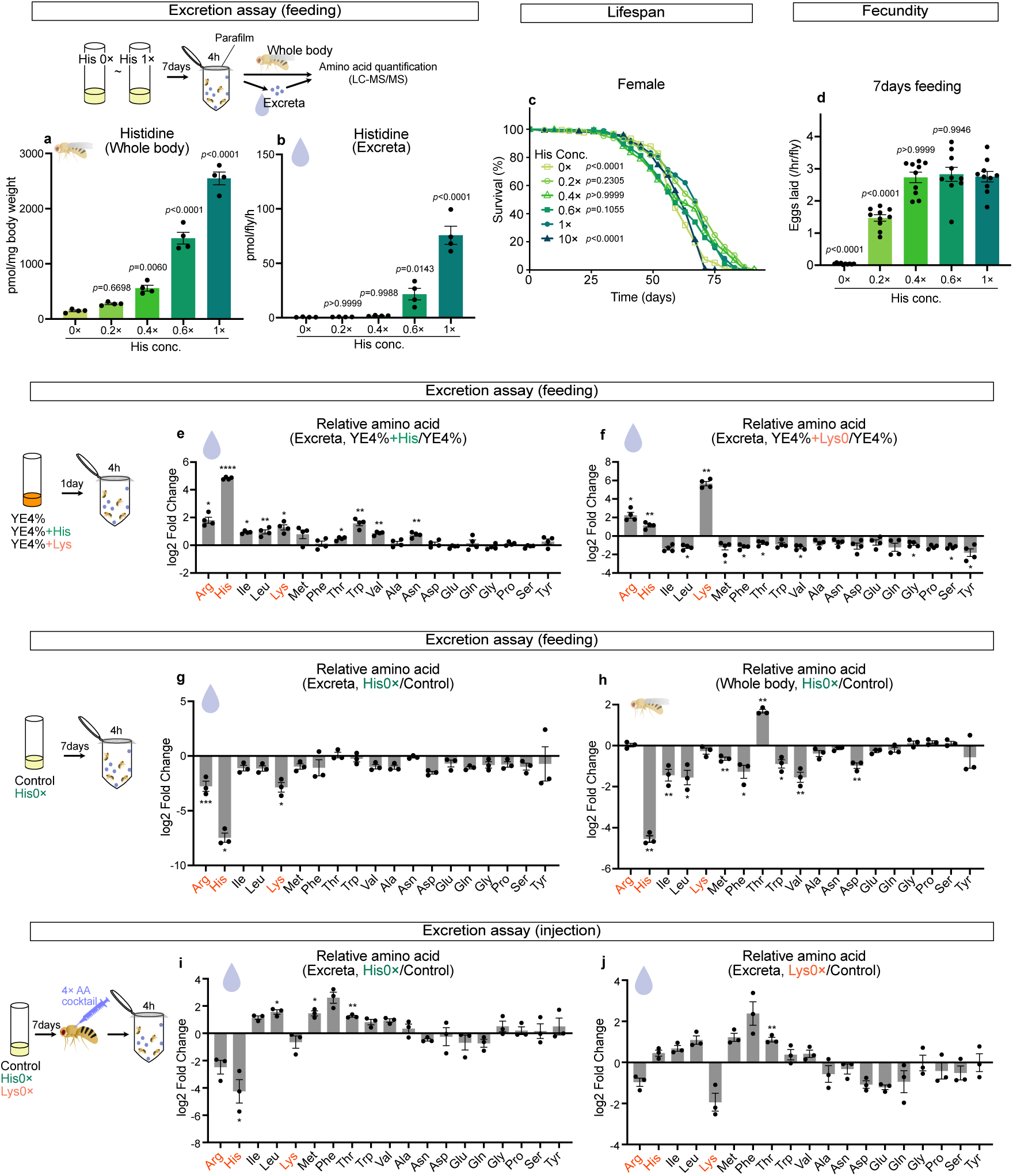
Excretion changes in response to alterations in individual dietary amino acid levels. **a, b** Quantification of His in whole bodies (**a**) and excreta of female CS flies reared on synthetic diets containing His at indicated concentrations for 7 days. Flies were placed in tubes for 4 h to collect excreta. n=4. Statistics: one-way ANOVA with Šidák’s multiple comparisons. **c** Lifespan of female CS flies reared on synthetic diets containing His at indicated concentrations. n=180 for His 0×, n=190 for His 0.2×, n=180 for His 0.4×, n=182 for His 0.6×, n=186 for His 1×, and n=184 for His 10×. Statistics: log-rank test. **d** Fecundity of female CS flies reared on synthetic diets containing His at indicated concentrations for 7 days. n=10. Statistics: one-way ANOVA with Šidák’s multiple comparisons. **e, f** Log2 fold changes in individual AA levels in excreta of female CS flies fed 4% YE diet supplemented with His (**e**) or Lys (**f**) to levels equivalent to those in a 12% YE diet for 1 day. Flies were placed in tubes for 4 h to collect excreta. n=3. **g, h** Log2 fold changes in individual AA levels in excreta of female CS flies reared on synthetic diets lacking either His (**g**) or Lys (**h**) for 7 days, normalised to the control. After injection with an AA cocktail, flies were placed in tubes for 4 h to collect excreta. n=3. **i, j** Log2 fold changes in individual AA levels in excreta of female CS flies reared on synthetic diets lacking either His (**g**) or Lys (**h**) for 7 days, normalised to the control. After injection with an AA cocktail, flies were placed in tubes for 4 h to collect excreta. n=3. *, *p*_adj_ <0.05, **, *p*_adj_ <0.01, ***, *p*_adj_ <0.001, ****, *p*_adj_ <0.0001. Each graph shows the mean ± SEM.

**Extended Data Fig. 8.**
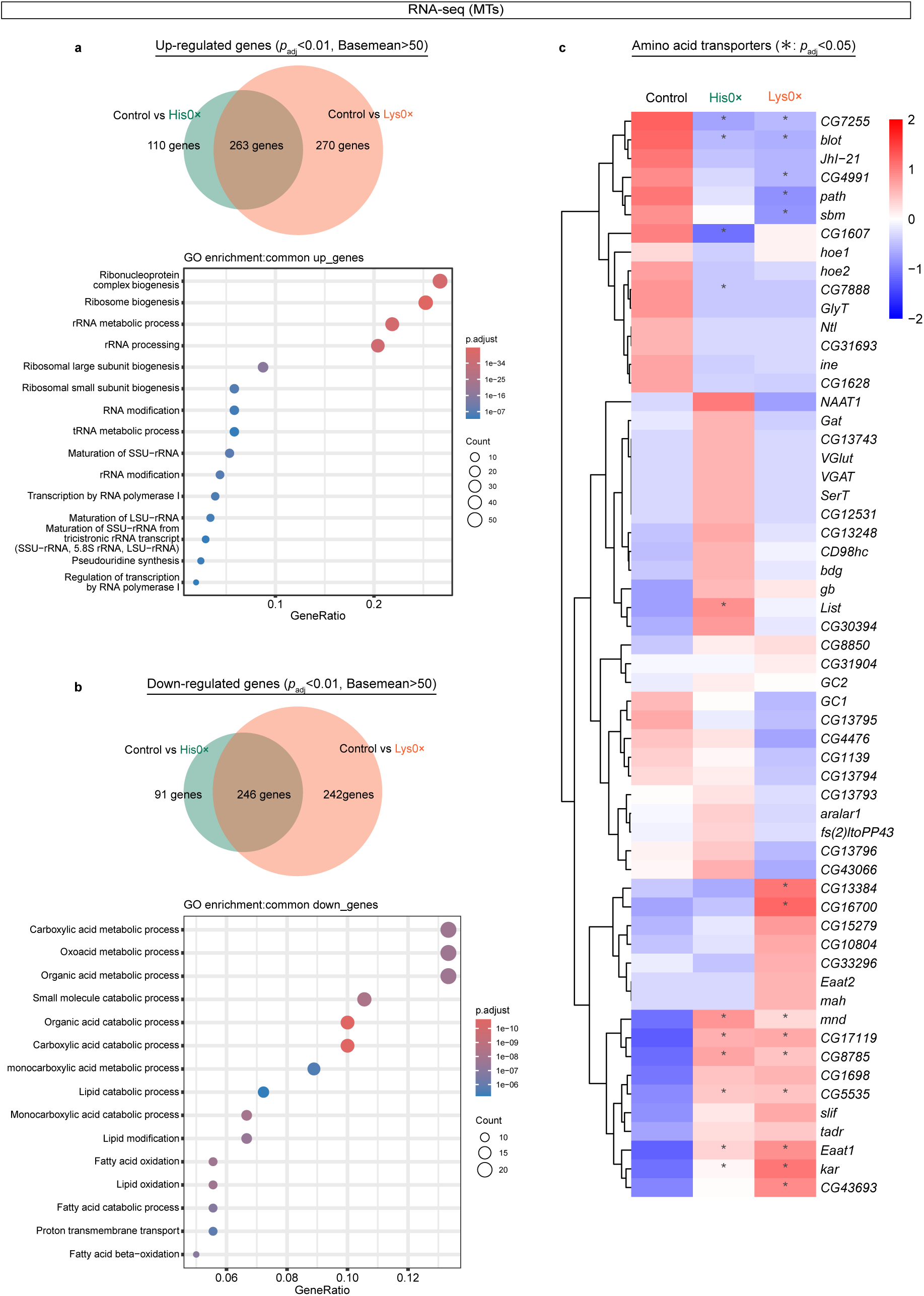
Transcriptomic analysis of Malpighian tubules from female CS flies fed His or Lys deficient synthetic diets for three days. **a, b** Venn diagrams illustrating the numbers of genes significantly upregulated (*p*_adj_<0.01, Basemean>50, Log2FoldChange>0) (**a**) or downregulated (*p*_adj_<0.01, Basemean>50, Log2FoldChange<0) (**b**) under His or Lys depletion (upper panels), along with dot plots showing the top 15 significantly enriched GO terms for genes commonly altered in both conditions (lower panels). **c** Heatmap showing expression changes of putative AA transporter genes under His or Lys depletion. *, *p*_adj_ <0.05, **, *p*_adj_ <0.01, ***, *p*_adj_ <0.001, ****, *p*_adj_ <0.0001.

**Extended Data Fig. 9.**
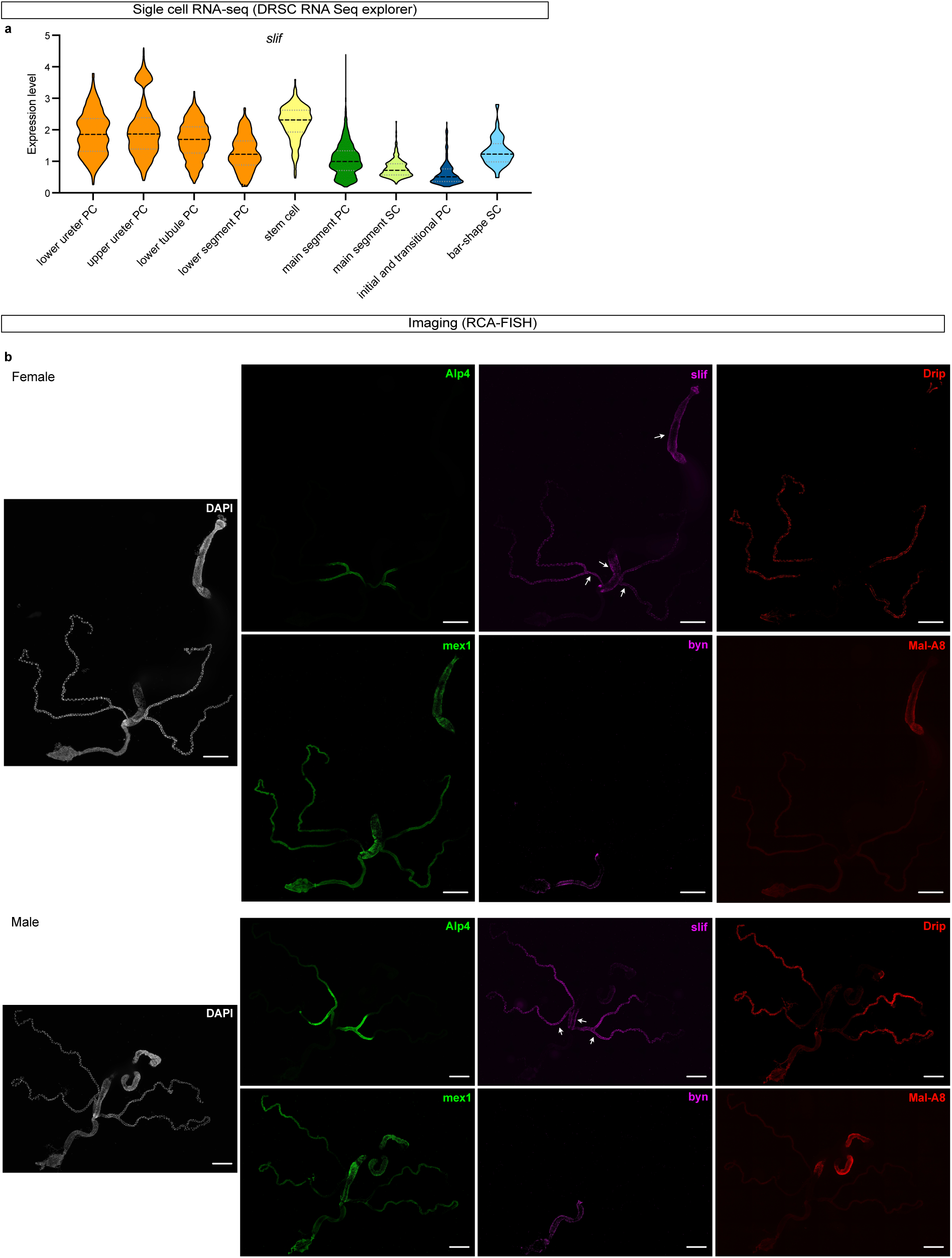
Localisation of *slimfast* in gut and MTs. **a** A violin plot of the expression level of *slif* in indicated cell types. PC, principal cell; SC, stellate cell. **b** Expression pattern of *slif* in the gut and MTs of 1-week-old female and male CS flies. White arrows in the *slif* panel indicate expression in the R5 region of the midgut and in the lower tubule. Scale bar: 500 µm.

**Extended Data Fig. 10.**
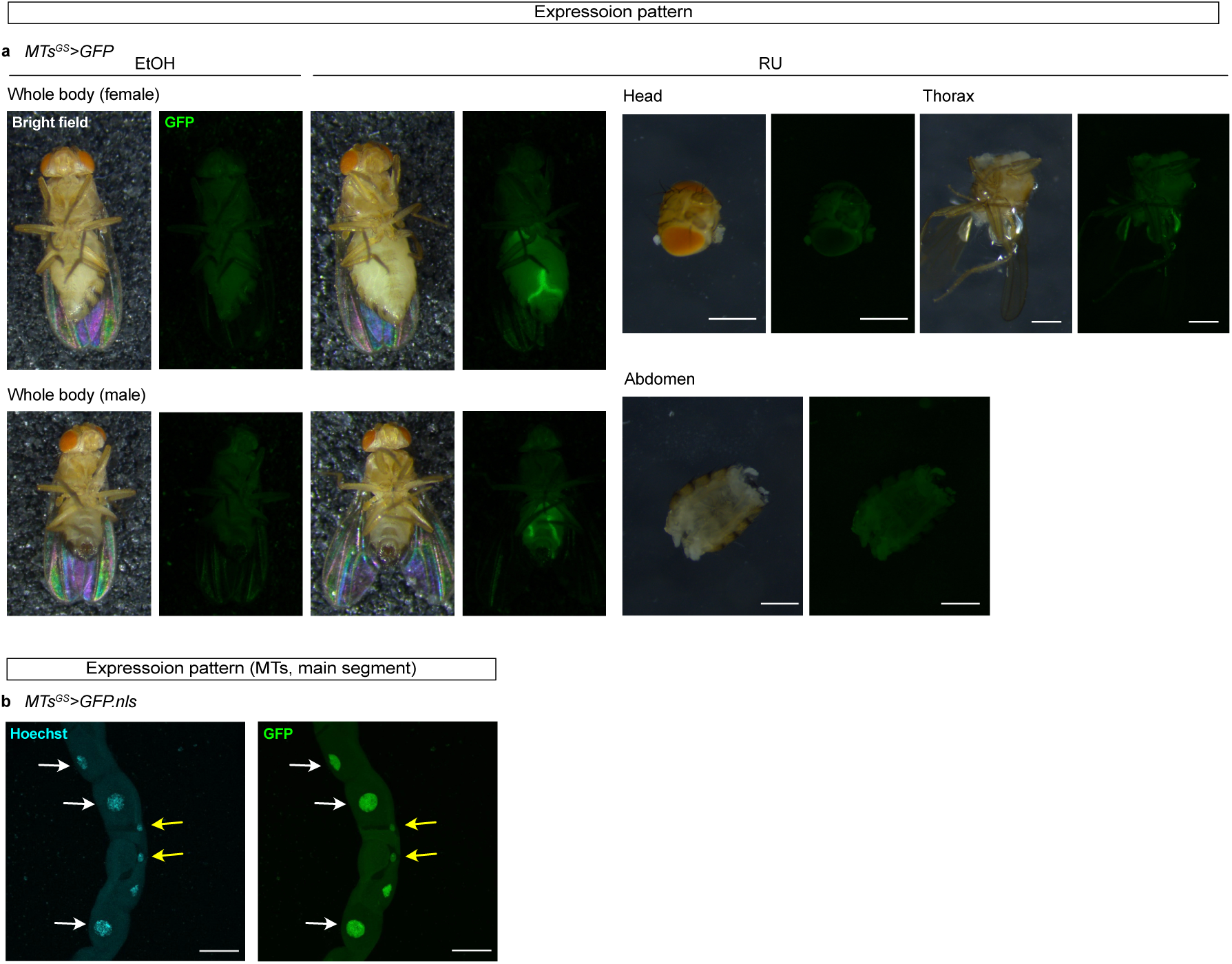
The expression pattern of *MTs^GS^* driver. **a** GFP expression of *MTs^GS^>UAS-GFP* in whole body and each tissue. Flies were fed a diet containing EtOH (control) 50 μM RU486 for 3 days. Green, GFP signal. Scale bar: 500 µm. **b** GFP expression of *MTs^GS^>UAS-GFP.nls* in the main segment of MTs. Flies were fed a diet containing 200 μM RU486 for 3 days. Cyan signals indicate Hoechst 33342 signal (nucleus) and green signals indicate GFP. White allows show principal cells and yellow allows show stellate cells with GFP signal. Scale bar: 50 µm.

